# Conserved developmental rewiring of the TCR signalosome drives tolerance in innate-like lymphocytes

**DOI:** 10.1101/2023.09.01.555859

**Authors:** Amanpreet Singh Chawla, Harriet J. Watt, Stefan A Schattgen, Neema Skariah, Irene Saha, Kathrynne A. Warrick, Masahito Ogawa, Jessica Strid, Frederic Lamoliatte, Alastair Copland, Sara Pryde, Elena Knatko, Kasper D. Rasmussen, Kazu Kikuchi, Paul G Thomas, Chandrashekar Pasare, David Bending, Mahima Swamy

## Abstract

Natural intraepithelial T lymphocytes (T-IELs) are innate-like, intestine-resident T cells essential for gut homeostasis. These cells express self-reactive T cell antigen receptors (TCRs) due to thymic agonist selection, but they do not cause autoimmunity. The mechanism underlying natural T-IELs tolerance in the gut is unclear. Using TCR reporter mouse models and phosphoproteomics, we demonstrate that TCR signaling is intrinsically suppressed in natural T-IELs. We discover that this suppression occurs post-selection in the thymus through altered expression of TCR signalosome components, a mechanism we term RePrESS (Rewiring of Proximal Elements of TCR Signalosome for Suppression). RePrESS is evolutionarily conserved and also found in autoreactive innate-like T cells from skin, breast and prostate. In coeliac disease, tolerance breakdown is associated with loss of RePrESS in natural T-IELs. Our findings reveal a distinct, conserved mechanism of tolerance involving TCR signaling rewiring, with implications for understanding barrier tissue homeostasis and autoimmune disease.

**One sentence summary:** Autoreactive innate-like T lymphocytes are developmentally suppressed by rewiring of the T cell antigen receptor signaling pathway to maintain tolerance.

## Introduction

Immune tolerance is critical for preventing autoreactivity and maintaining homeostasis despite continuous exposure to self-antigens, commensals and pathogens. Thymic selection eliminates most self-reactive T cells via negative selection, yet certain self-reactive T cells undergo a process known as agonist selection. In agonist selection, thymocytes with T cell antigen receptors (TCR) that recognize self-antigens with relatively high affinity receive strong TCR signals that drive their differentiation into specialized lineages, such as natural regulatory T cells (natural Tregs), invariant natural killer T (iNKT) cells, and natural intestinal intraepithelial T cells (natural T-IELs) ^1^. These cells have important roles in immune regulation and preventing autoimmunity, but it is unclear how these self-reactive T cells do not themselves trigger autoimmunity.

Natural T-IELs are particularly intriguing. Residing within the epithelial layer of the small intestine, T-IELs act as sentinels that mediate epithelial repair to maintain the intestinal barrier and immune surveillance to respond to pathogens while remaining tolerant of a complex array of dietary antigens, microbiota-derived molecules, and self-antigens ^2^. T-IEL represent one of the most abundant and evolutionarily ancient lymphocyte populations in the body. They are highly adapted to the intestinal environment, share many of the features of cytotoxic T cells, yet are tightly controlled in an activated-yet-resting state that limits their activation in homeostasis^3–5^. This tight control of the cytotoxic potential of T-IEL is clearly important, as in coeliac disease, a gluten-induced autoimmune disease, the damage of the intestine is driven by IFNγ- and GzmB-expressing T-IEL that are highly expanded in the diseased intestinal epithelium ^6^. Yet T-IEL also play important immunoregulatory roles, and can inhibit experimentally-induced colitis^7,8^. Understanding how these cells remain tolerant while retaining effector capacity is of fundamental immunological importance, with implications for mucosal homeostasis, autoimmune diseases such as celiac disease and Crohn’s.

T-IELs are classified into two main populations based on their ontogeny: induced and natural^9^. Induced T-IELs express αβ TCRs with CD4 or CD8αβ coreceptors, undergoing conventional thymic selection before exiting as naïve CD4+ or CD8+ T cells ^9^. In the periphery, induced T-IELs encounter antigen from commensals or diet, upregulate gut-homing molecules, and migrate into the intestinal epithelium. By contrast, natural T-IELs express either αβ or γδ TCRs and are defined by their expression of the CD8αα coreceptor, which is upregulated upon entry into the intestinal epithelium. Natural γδ T-IELs develop from CD4^–^ CD8^–^ double-negative (DN) thymocytes, commit to γδ T cell development, exit the thymus in a naïve state, and mature in the intestinal epithelium after activation in gut-associated lymphoid tissues. The γδ TCR expressed by a subset of γδT-IEL (Vγ7+) binds to butyrophilin-like (Btnl) molecules expressed by intestinal epithelial cells, suggesting cognate antigen recognition even under homeostatic conditions^10^. Natural αβ T-IELs follow a conventional T cell developmental trajectory through the Cd4+ CD8+ double positive (DP) stage until they receive strong agonist signals in the thymus. Natural αβT-IEL precursors express self-reactive TCRs that elicit stronger signals than those required for negative selection^11^, driving these cells to differentiate into T-IEL progenitors marked by downregulation of CD4 and CD8 and sequential upregulation of CD5, PD-1, and CD122 ^12^. Mature natural αβT-IEL precursors subsequently acquire gut-homing molecules in the thymus before migrating to the intestinal epithelium.

Despite expressing autoreactive TCRs, natural T-IELs exhibit considerable tolerance to self-ligands in the gut. In TCR-transgenic mice where the known TCR ligand is expressed in the thymus, agonist selection positively selects for natural αβT-IEL differentiation, but their TCR becomes hyporesponsive to the cognate ligand^13–15^. Even when the TCR ligand was expressed in the intestine, no autoreactivity was seen^15^. Moreover, viral infections that often lead to loss of T cell tolerance to self-antigens, do not affect the self-tolerance of natural αβT-IEL^15^. This remarkable balance between autoreactivity and functional competence has made natural T-IELs a central but unresolved model for studying peripheral tolerance.

Conventional TCR signaling pathways are well characterized: TCR engagement triggers phosphorylation of immunoreceptor tyrosine-based activation motifs (ITAMs) on CD3 chains by Lck, which recruits ZAP-70 and initiates downstream signaling through the phosphorylation of the adaptor protein LAT ^16,17^. Phosphorylated LAT forms a scaffold for critical downstream effectors, including SLP-76, PLCγ1, and Itk, leading to calcium mobilization, MAPK activation, and full T cell activation. Whether natural T-IELs retain these pathways is unclear. Notably, natural T-IELs display reduced LAT expression and incorporation of the FcεR1γ chain in place of CD3ζ in their TCR complex^18^. Despite these observations, TCR signaling in natural T-IELs has not been characterized.

Here, we uncover the molecular basis of tolerance in natural T-IELs, revealing a mechanism that we term RePrESS (Rewiring of Proximal Elements of TCR Signalosome for Suppression). Using phosphoproteomics, we demonstrate that natural T-IELs exhibit extensive rewiring of their TCR signaling pathways compared to conventional T cells and induced T-IELs. Strikingly, natural T-IELs are intrinsically refractory to TCR stimulation, characterized by diminished phosphorylation of key signaling intermediates. This hyporesponsiveness is established during thymic development and is conserved in zebrafish, mice, pigs, and humans. Likewise, rewiring of the TCR signalosome is observed in mammary, prostate and skin innate-like T-IELs, and is partially conserved in other agonist-selected T cells. Notably, in celiac disease, the rewiring of TCR signaling is reversed in natural T-IELs, thus leading to a breakdown of tolerance, and autoimmune pathology. Therefore, our findings reveal a previously unknown, broadly applicable, cell autonomous mechanism of innate-like T cell tolerance, which has substantial implications for understanding peripheral tolerance and immune dysregulation in barrier tissues in disease.

## Results

### TCR signal transduction is suppressed in autoreactive T-IEL

It has previously been suggested that natural T-IELs are hyporesponsive to TCR stimulation in vitro^19,20^. To test whether natural T-IELs are refractory to TCR stimuli *in vivo*, we employed Nur77-Tempo mice^21^, a TCR signaling reporter model that tracks signaling in a temporal and graded manner using the largely NFAT-independent gene *Nr4a1* (which encodes Nur77) (**Fig. 1a**)^22^. In this transgenic model a fluorescent timer protein is expressed under the control of the TCR inducible gene *Nr4a1*^22^. Initially upon translation, the Timer protein has blue fluorescence before undergoing irreversible maturation to a red fluorescent form with an approximate blue half-life of 4 hours and red half-life of >5 days (**Fig. 1a**). This reporter is very sensitive and specific to antigen receptor triggering due to its dependence on multiple signaling pathways downstream of the TCR ^21^, and can therefore reveal whether the TCR has been recently activated in cells. Intraperitoneal anti-CD3 injection in Nur77-Tempo mice elicited Nur77-blue signal in 80-90% of conventional CD8+ T cells, but only in <10% αβ and in ∼25% γδ natural T-IELs (**Fig. 1b**). Moreover, the few blue-positive natural T-IELs had very low blue intensities, indicating overall TCR hyporesponsiveness (**Fig. 1b**). Further, both natural T-IEL populations produce less IFNγ, and show reduced expression of CD107, a marker of degranulation, compared to induced T-IEL when stimulated with plate-bound anti-CD3 directly *ex-vivo* (**Fig. 1c**).

**Figure 1.**
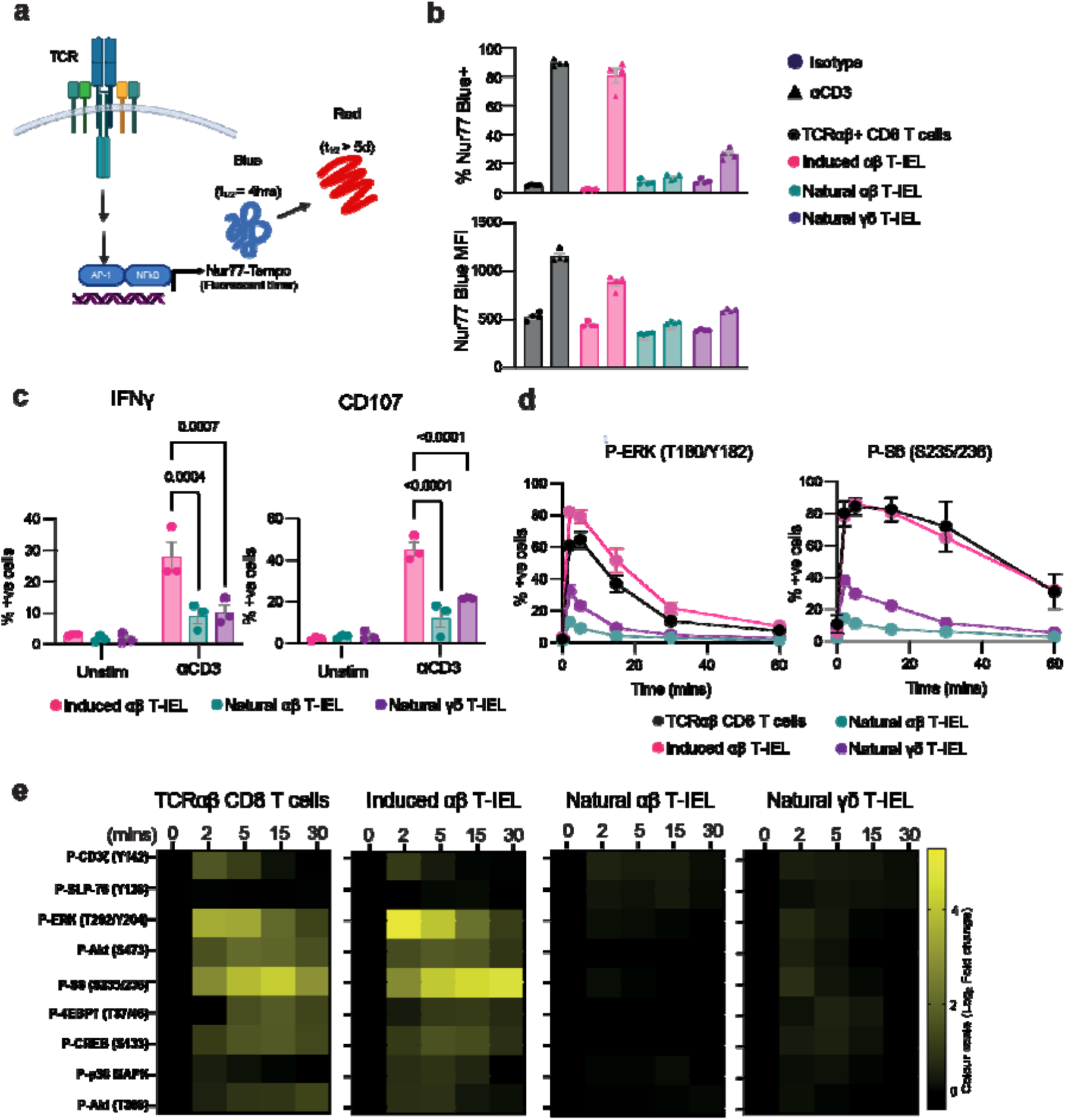
TCR signaling is suppressed in autoreactive intestinal natural αβT-IEL. (a) Schematic representation of the Nur77-Tempo reporter system. This transgenic *Nr4a1*-dependent fluorescent timer model tracks TCR activation dependent on multiple signaling inputs. Blue fluorescence (t₁/₂ ≈ 4 hours) indicates recent TCR activation, and red fluorescence (t₁/₂ > 5 days) reflects historical TCR engagement. Created in BioRender. (b) The percentage (top) and MFI (bottom) of Nur77-blue+ cells was quantified 4 hours after intraperitoneal injection of 50 μg anti-CD3 or isotype control antibody in Nur77-Tempo reporter mice. Data represents mean ± SEM from four biological replicates. (c) The percentage of T-IEL subsets positive for IFNγ and CD107 was quantified after ex vivo stimulation with plate-bound anti-CD3 (3 μg/ml) for 4 hours. Data show the mean ± SEM from three biological replicates, representative of at least 3 independent experiments. (d) The percentages of indicated T cell subsets positive for phosphorylation of ERK1/2 (T180/Y182) and ribosomal protein S6 (S235/S236) were measured following *ex vivo* stimulation with anti-CD3 (30 μg/ml) crosslinked with anti-Hamster (5 μg/ml). Data shows the mean ± SEM from five independent experiments. (e) Heatmaps depicting the activation of key TCR signaling molecules in conventional CD8+ T cells and T-IEL subsets. Log₂ fold changes in mean fluorescence intensity (MFI) of phosphorylated proteins were measured in conventional CD8 T cells and T-IEL subsets following anti-CD3 (30 μg/ml) stimulation crosslinked with anti-Hamster (5 μg/ml). Fold changes are calculated relative to unstimulated conditions (Time 0). Data represents the mean from at least three independent biological replicates. Statistical significance was determined using ANOVA with Tukey’s post hoc correction. Non-significant values are not shown.

Since natural T-IEL responded poorly to TCR stimulation, we next assessed the TCR signaling pathway. To allow us to stimulate and compare T-IELs with conventional TCRαβ CD8+ T cells in identical conditions, we modified the fluorescent cell barcoding technique ^23^ to differentially label T-IELs and conventional CD8+ T cells. T-IELs were labelled with an amine-reactive dye and combined with unstained conventional CD8+ T cells from lymph nodes, allowing these cells to be stimulated in the same tube as T-IELs and subsequently differentiated from T-IELs during flow analysis **(Suppl. Fig. 1a)**. The cells in the various stimulation conditions were then subject to fluorescent cell barcoding prior to phospho-flow cytometry. The specificities of phospho-antibodies were verified using inhibitor-treated cells **(Suppl. Fig. 1b)**. Both conventional CD8+ T cells and the induced T-IEL subset showed strong responses to TCR stimulation, with >60% of cells positive for phosphorylation of ERK1/2 and the ribosomal protein S6 (**Fig. 1d**). By contrast, agonist-selected natural αβ T-IELs remained largely unresponsive, with fewer than 15% exhibiting phosphorylation, while natural γδ T-IELs showed similarly muted activation (<40%). This significant reduction in TCR-induced activation of natural compared with induced T-IEL subsets is evident in the heatmap showing the log_2_ fold change in median fluorescence intensity (MFI) for multiple TCR signaling proteins (**Fig. 1e**). Thus, decreased phosphorylation of key signaling nodes downstream of TCR engagement in natural T-IELs also translated into reduced functional responses, particularly of the agonist-selected natural αβ T-IEL, confirming that natural T-IELs are refractory to TCR stimulation.

### Tolerance in natural T-IEL is independent of Tregs, anergy or chronic stimulation

Regulatory T cells (Tregs) are widely recognized as the central enforcers of immune tolerance, critical for suppressing self-reactive T cells and preventing autoimmunity. To investigate whether Tregs are essential for restraining self-reactive natural T-IELs and preventing intestinal autoimmunity, we employed the Foxp3-DTR mouse model ^24^, which enables specific ablation of Tregs upon diphtheria toxin (DT) administration. As expected, Treg depletion triggered widespread activation of systemic conventional CD8 T cells, characterized by a robust increase in CD44 and effector molecule expression, including TNF, IFNγ, and granzyme A in splenic conventional CD8+ T cells (**Fig. 2a**). Surprisingly, natural αβ T-IEL and γδ T-IEL in the small intestine remained unaffected. These cells showed no upregulation of activation markers such as CD44 or granzyme A, nor did they produce elevated levels of TNF or IFNγ (**Fig. 1a** and **Suppl. Fig. 1**). Histological analysis further confirmed the absence of inflammation or epithelial damage in the small intestine, reinforcing that natural T-IELs remain quiescent and are not reliant on Treg-mediated suppression to maintain their tolerized state (**Fig. 1b**). A similar effect was seen in *foxp3a^-/-^* zebrafish ^25^, which harbor IEL-like cells that express granzymes in their intestines^26^. We found that Treg deficiency in *foxp3a-/-* zebrafish triggered robust inflammation in peripheral tissues, including the gills, but the intestines remained completely unaffected, with no signs of inflammation or immune cell infiltration (**Suppl. Fig. 1b,c**). Thus, gut T-IEL tolerance is evolutionarily ancient and operates independently of Tregs, pointing to a reliance on cell-intrinsic mechanisms for their regulation.

**Figure 2.**
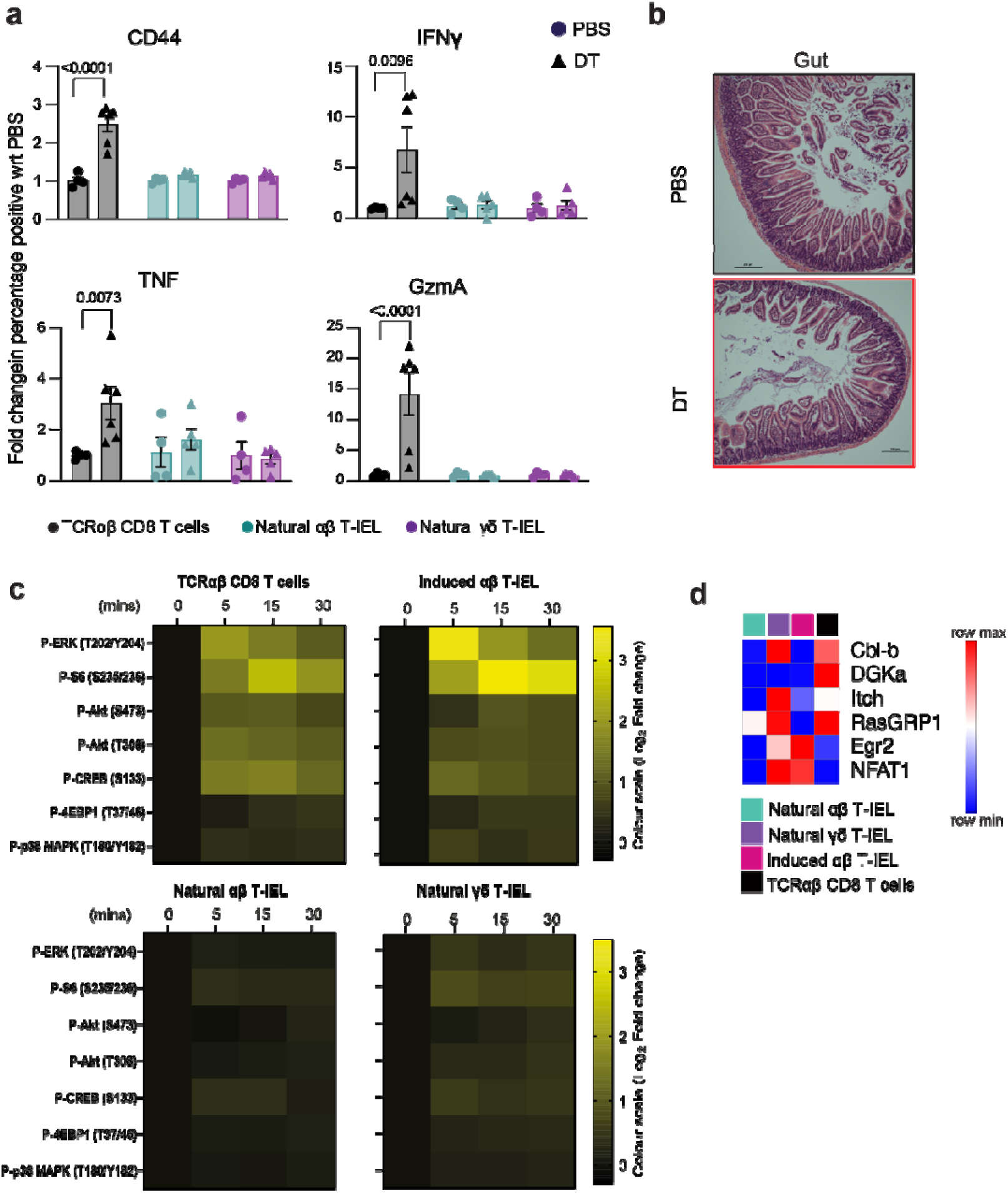
T-IEL tolerance is independent of regulatory T cells chronic stimulation or anergy. Foxp3-DTR mice were treated with PBS or diphtheria toxin (DT) on days 0, 3 and (**a**) immune activation was assessed on day 5 post-first DT administration. Flow cytometric analysis quantified the fold change in immune activation markers (CD44), inflammatory cytokines (IFNγ, TNF), and cytotoxic markers (granzyme A, GzmA) in conventional CD8α+ T cells from the spleen and the intestinal intraepithelial lymphocyte compartment (T-IEL). Data represents the mean ± SEM from at least four biological replicates. (**b**) Histological analysis of small intestine inflammation at day 5 post-DT treatment. Representative H&E-stained images of the small intestine from PBS- and DT-treated mice illustrate the presence or absence of inflammatory pathology. (**c**) Heatmap quantifying anti-CD3-induced TCR signaling in purified conventional CD8+ T cells and T-IEL subsets following four days of culture with IL-15. Log_2_ fold changes are calculated relative to no anti-CD3 conditions (time 0). Data depict mean from four biological replicates. (**d**) Row-normalized heatmap of protein copy numbers of selected proteins important in maintaining anergy, in T-IEL subsets from published proteomics data ^4^.

One potential explanation for their lack of response to TCR triggering might be that natural T-IELs experience chronic TCR stimulation in the gut, making them refractory to further signaling. If this is the case, removing natural T-IELs from the gut should restore their responsiveness. To test this, we purified T-IELs from WT mice and cultured them with IL-15 for four days to allow the cells sufficient time in a ligand-free environment before stimulating them with anti-CD3. However, even in the absence of potential cognate ligands, and outside the immunosuppressive environment of the gut, natural T-IEL remained unresponsive to TCR stimulation (**Fig. 2c**).

As natural T-IEL do not express CD28 ^18^, we next checked whether natural T-IELs are anergic due to lack of co-stimulation. However, natural T-IEL are not anergic as they do not express markers of anergy ^27^ such as high expression of Cbl-b, DGKα or Itch E3 ligase that downregulate TCR signaling, based on previously published proteomic comparisons of T-IEL and conventional CD8 T cells ^4^ (**Fig. 2d**). Collectively, these results establish that the hyporesponsive nature of natural T-IELs in the intestine is not driven by exhaustion from chronic TCR stimulation or anergy due to lack of co-stimulation but likely reflects a distinct cell-autonomous tolerance mechanism.

### Rewiring of proximal TCR signaling in natural T-IEL

To investigate alternative tolerance mechanisms, we examined the TCR signaling pathways in natural T-IELs. We utilized phorbol myristate acetate (PMA) and ionomycin, which bypass TCR ligation and directly activate key downstream pathways. PMA activates protein kinase C (PKC) ^28^; while ionomycin is a calcium ionophore that triggers calcium flux ^29^, thus, mimicking TCR signaling by targeting the pathways downstream of phospholipase-Cγ (PLCγ) and PKC. Remarkably, stimulation with PMA and ionomycin induced robust phosphorylation of ERK1/2 and S6 in 100% of T-IEL subsets **(Fig. 3a)**. Additionally, phosphorylation of p38 MAPK and CREB in T-IELs was comparable to that observed in induced T-IEL (**Fig. 3c and Suppl. Fig. 3a-b)**. These findings confirm that the signaling pathways downstream of PLCγ and PKC remain intact in natural T-IELs, induced T-IELs, and conventional CD8 T cells.

**Figure 3.**
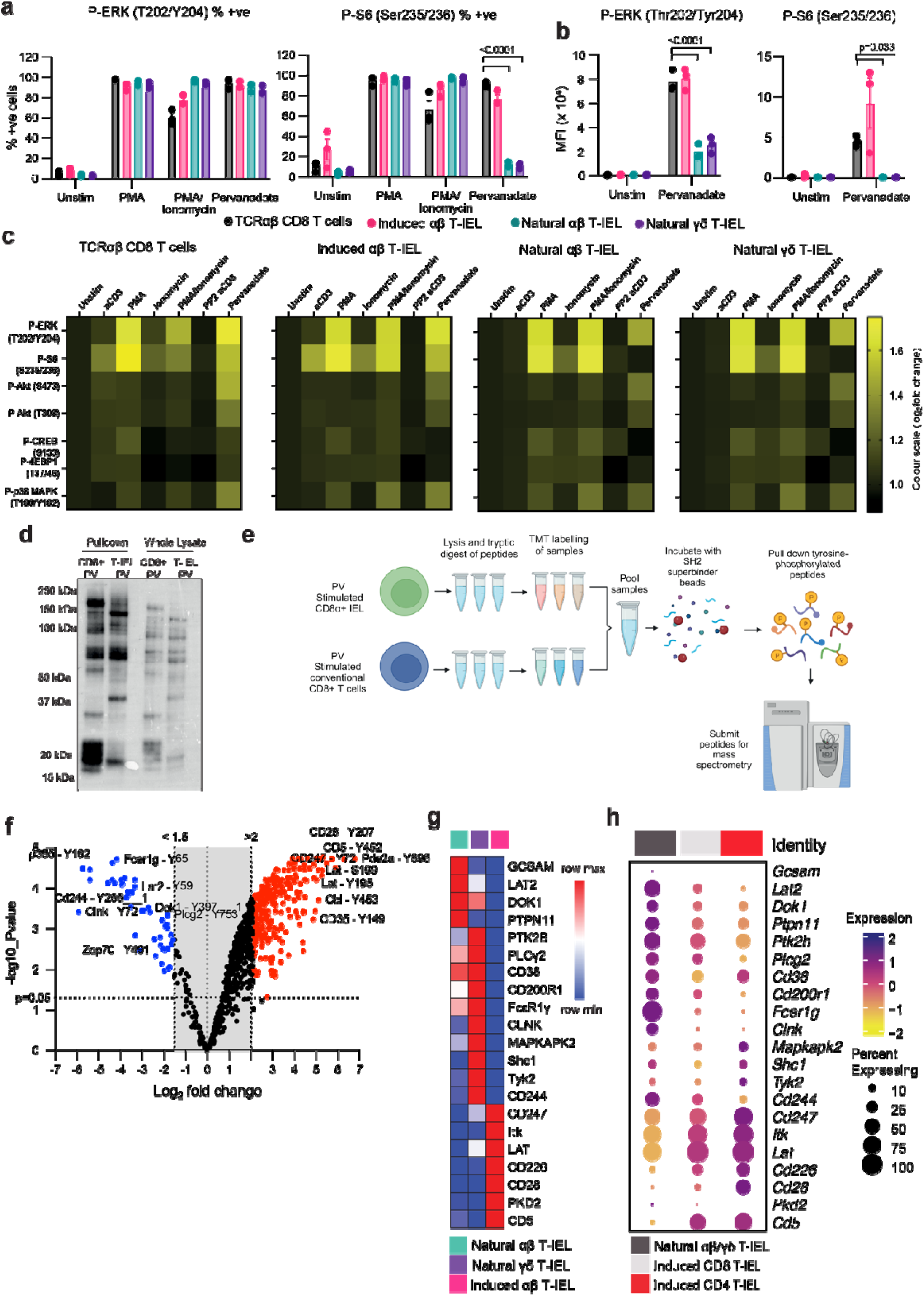
Phosphoproteomics reveals a rewired TCR signalosome signature in natural T-IEL. (**a, b**) Bar plots showing (**a**) the percentage of cells positive and (**b**) the mean fluorescence intensity (MFI) of phosphorylation of key signaling proteins following stimulation with PMA (100 ng/ml), Ionomycin (1 μg/ml), or pervanadate (10 μM). Data represent mean ± SEM from three biological replicates. (**c**) Heatmaps depict the log₂ fold change in MFI of phospho-proteins across T-IEL subsets and conventional CD8α+ T cells, under the same stimulation conditions as in (a) and anti-CD3 as in Fig. 2. 10 μM PP2 was added to the indicated condition and incubated for 1 hour prior to stimulation. Log_2_ fold change is calculated relative to the unstimulated condition. Data represents mean from three biological replicates. (**d**) Phospho-tyrosine immunoblot of lysates from pooled CD8α+ αβ and γδ natural T-IEL or conventional CD8α+ T cells stimulated with pervanadate (10 µM) for 5 minutes and enriched for phosphotyrosine-containing proteins using SH2 superbinder beads. (**e**) Experimental procedure for the preparation of samples for phospho-proteomics. (**f**) Volcano plot of log_2_ fold change and log_10_ p-value of the phospho-sites identified in natural T-IEL vs. conventional CD8+ T cells. Data are from three biological replicates. (**g**) Row-normalized heatmap of protein copy numbers from T-IEL subsets of proteins identified as differentially phosphorylated in (f), from published proteomics data ^4^. (**h**) Bubble plots indicating gene expression levels and frequency of cells expressing TCR signalosome genes within murine T-IEL clusters derived from single-cell RNA-sequencing data ^33^.

Given the intact PLCγ- and PKC-dependent pathways, we next assessed the functionality of TCR-proximal signaling proteins. To do so, we used pervanadate, a pan-tyrosine phosphatase inhibitor that mimics TCR stimulation by blocking TCR-proximal phosphatases, such as CD45, and activating proximal kinases such as Lck and ZAP70 (32). While pervanadate induced phosphorylation of ERK and Akt across all T-IEL subsets **(Fig. 4a-c, Suppl. Fig. 4)**, it failed to drive phosphorylation of key downstream signaling proteins such as S6 and CREB in natural T-IEL subsets. Furthermore, the magnitude of ERK and Akt phosphorylation was significantly lower in natural T-IELs compared to conventional CD8 T cells. These results indicate a disruption in TCR signal transduction in natural T-IEL that likely occurs at the TCR-proximal stage, as PMA (which bypasses TCR proximal signaling) restored phosphorylation of ERK1/2, S6, and CREB, while pervanadate could not. Therefore, while downstream signaling pathways are intact, proximal TCR signaling in natural T-IELs is impaired.

**Figure 4.**
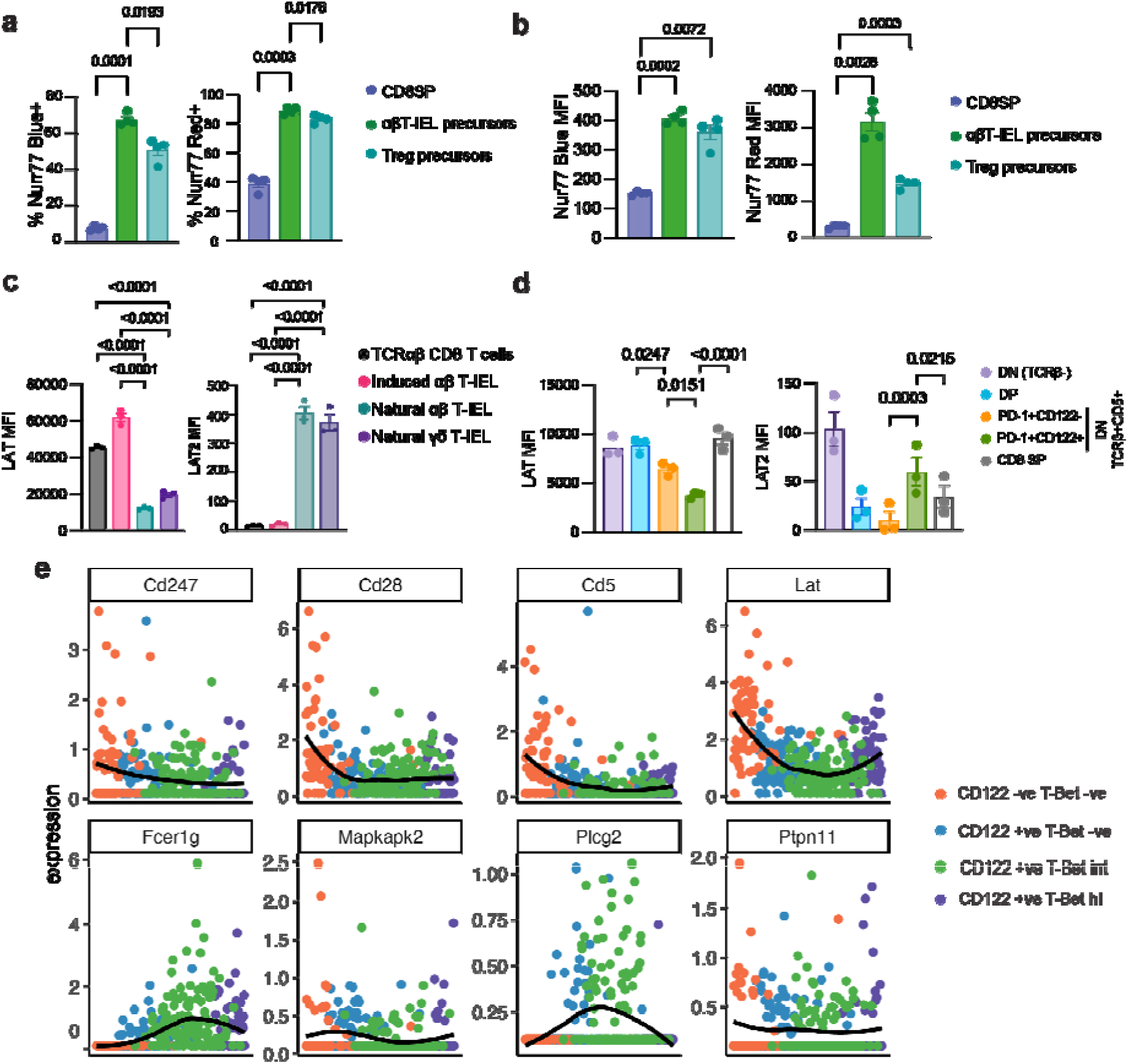
The TCR signalosome is rewired during thymic selection of Natural T-IEL. (a, b) Bar graphs show (a) percentage positive and (b) MFI of Nur77-blue and Nur77-red in post-selection thymic precursors of natural αβ T-IEL, Treg precursors, and conventional CD8+ single-positive (CD8SP) thymocytes in Nur77-Tempo reporter mice at steady state. Gating strategy shown in Suppl. Fig. 6a-b. Data represents mean ± SEM from four biological replicates. (**c, d**) Flow cytometric analysis of LAT and LAT2 expression in (c) mature intestinal T-IEL subsets compared to conventional CD8+ T cells, and in (d) thymocyte populations, including double-negative (DN) thymocytes (CD4⁻CD8⁻ TCRβ^–^), double positive (DP) thymocytes (CD4^+^CD8^+^TCRβ^lo^), IEL precursors (CD4⁻CD8⁻TCRβ⁺CD5⁺PD-1^+^CD122^–^ or the more mature CD4⁻CD8⁻TCRβ⁺CD5⁺PD-1⁺CD122⁺), and conventional CD8 T cell precursors (CD8 SP). Data in c and d are mean ± SEM from three biological replicates. (e) Expression profiles of selected TCR signalosome genes along the predicted IEL precursor differentiation trajectory were analyzed from single-cell RNA sequencing data from (Hummel et al., Mucosal Immunology, 2020)^34^. Statistical significance (a-d) was determined using ANOVA with Tukey’s post hoc correction.

It is possible that natural T-IELs engage noncanonical TCR signaling pathways that are distinct from those utilized by conventional CD8 T cells. Immunoblotting of phosphotyrosine-containing proteins enriched from natural T-IEL and CD8 T cells after pervanadate stimulation, revealed global differences in phosphorylated proteins between natural T-IELs and conventional CD8 T cells **(Fig. 3d)**. Therefore, we sought to identify the molecular bases of this distinct signaling phenotype. We conducted an unbiased phosphoproteomic analysis of tyrosine phosphorylation events in pervanadate-stimulated natural T-IELs and conventional CD8 T cells. Approximately 10 million natural T-IELs were pooled from the intestines of 10 WT C57Bl/6 mice per replicate, sorted for CD8α+ CD8β-CD4-, then stimulated for 5 minutes with pervanadate to amplify TCR-associated phosphorylation. The natural T-IEL population used for phospho-proteomic analyses consisted of both αβ and γδ subsets, as sufficient cell numbers could not be obtained from the individual populations. An equivalent number of conventional CD8 T cells were isolated from lymph nodes and spleens, sorted and stimulated with pervanadate **(Suppl. Fig. 4a)**. Pervanadate-stimulated samples were lysed, digested and labeled using Tandem Mass tagging, then pooled **(Fig. 3e)**. Tyrosine-phosphorylated proteins were pulled down using high affinity SH2 superbinder beads that specifically bind to phospho-tyrosine ^31,32^. In total we identified 819 phospho-tyrosine containing phospho-peptides in the natural T-IELs and conventional CD8 T cells. Principal component analyses revealed that phosphopeptides from natural T-IELs and conventional CD8 T cells cluster away from each other and most of the difference is explained by the differences in cell type **(Suppl. Fig. 4b)**. Clustering the significantly regulated phospho-sites in a heatmap further corroborates large differences between natural T-IELs and lymph node samples, while showing good correlation between biological replicates **(Suppl. Fig. 4c)**.

Further investigation of the differentially phosphorylated peptides reveals overwhelmingly more proteins are phosphorylated in conventional CD8 T cells (570 phospho-sites) as compared to 60 phospho-sites significantly enriched in the natural T-IEL population **(Fig. 3f)**. GO term analysis of the proteins that were significantly upregulated in CD8 T cells confirms that pervanadate is a good mimic of TCR signaling in conventional T cells, whereas in T-IELs, phosphoproteins were largely involved in negatively regulating immune system processes and those associated with myeloid and B cells **(Suppl. Fig. 4d, Suppl. tables 1 and 2)**. Intriguingly, many phospho-peptides enriched in conventional CD8 T cells belong to TCR signalosome proteins such as CD28, CD5 and LAT (Linker for activation of T cells) and TCR components CD3γ and CD3ζ (CD247), which our prior proteomics analyses ^4^ showed are either absent or expressed at much lower levels in T-IELs **(Fig. 3g)**. Conversely, peptides found highly phosphorylated in T-IELs belong to proteins that are exclusively expressed in natural T-IELs among T cells, including LAT2, GCSAM, PLCγ2, FcεR1γ, CLNK (**Fig. 3g)**. Based on the proteins that were both differentially phosphorylated in the current phosphoproteomics dataset and differentially expressed at the protein level, we curated a natural T-IEL protein signature **(Fig. 3g)** that we term RePrESS (**R**ewiring of **Pr**oximal **E**lements of the TCR **S**ignalosome for **S**uppression). We cross-referenced this signature with single-cell RNA-sequencing data from unstimulated WT mouse natural and induced T-IEL ^33^. We found that our curated RePreESS protein signature was conserved at the RNA level in natural T-IELs **(Fig. 3h)**. Of note, in scRNA-seq clustering, natural αβ and γδ T-IELs could not be clearly distinguished and thus are shown here together, supporting our pooling of the two natural T-IEL subsets for phosphoproteomics. The conservation of the natural T-IEL RePrESS signature at the transcript level confirms that the TCR signaling network in natural T-IELs is substantially rewired compared to conventional CD8 T cells and induced T-IEL. The differences appear to be driven largely by differential gene expression, resulting in a distinct signaling architecture that diverges significantly from classical TCR signaling pathways and is likely responsible for T-IEL tolerance.

### The TCR signalosome of natural **αβ**T-IELs is rewired during thymic maturation

Agonist selection requires strong TCR signaling, indicating that in thymic natural αβT-IEL precursors, TCR signaling must be intact. To explore TCR signaling in αβT-IEL precursors, we analyzed the thymus of Nur77-Tempo mice. As expected, natural αβT-IEL precursors (defined as CD4-CD8-CD25-CD1d-TCRβ+CD5+PD-1+, Suppl. Fig. 7a) were all highly positive for Nur77-blue signal, indicative of recent strong TCR stimulation during agonist selection **(Fig. 4a-b)**. Interestingly, natural αβT-IEL precursors were positive for both Nur77-blue and Nur77-red signals **(Suppl. Fig. 6b)** and displayed the highest Nur77-red signals among all thymocyte populations, surpassing even natural Tregs **(Fig. 4b)**. This observation suggested that natural T-IEL precursors undergo exceptionally strong and persistent TCR stimulation during thymic selection.

As TCR signaling is intact in αβT-IEL precursors, but not in mature intestinal IEL, we hypothesized that rewiring of the TCR signalosome occurs post-selection in the thymus. To test this hypothesis, we evaluated the expression of LAT and LAT2, two important adaptor proteins in antigen receptor signaling, using specific antibodies **(Suppl Fig. 6c-d).** We confirmed that LAT is more highly expressed in conventional CD8 T cells and induced T-IELs, whereas mature natural T-IELs express LAT2 **(Fig. 4c)**. We then compared the expression of LAT and LAT2 in early (TCRβ^+^ CD5^+^ PD-1^+^CD122^-^) and more mature (TCRβ^+^CD5^+^ PD-1^+^CD122^+^) post-selection thymic αβ T-IEL precursors cells versus conventional CD8 single positive (SP) precursors, and pre-selection double negative (DN, CD4-CD8-) and double positive (DP, CD4+CD8+) thymocytes. Strikingly, LAT expression was progressively downregulated, while LAT2 expression was upregulated, as natural T-IEL precursors matured along the thymic developmental trajectory **(Fig. 4d and Suppl Fig. 6e)**. These changes in LAT and LAT2 expression were unique to natural αβ T-IEL precursors and not observed in thymocytes that give rise to conventional CD8 T cell**s**.

Beyond LAT and LAT2, we sought to investigate whether other elements of the rewired TCR signalosome were similarly modulated in the thymus. Using published single-cell RNA sequencing data of mouse TCRαβ IEL precursors in the thymus ^34^, we performed trajectory analysis of all the genes in the RePreSS signature **(Fig. 4e and Suppl. Fig 6f)**. We observed downregulation of *Cd247* (CD3ζ), *Cd28, Cd5*, *Lat* and upregulation of RNAs including *Fcer1g*, *Plcg2*, *Tyk2* and *Ptpn11* (SHP2) along the precursor maturation pathway. These findings identify the thymus as the origin of the distinct molecular program that shapes the TCR signalosome and hence tolerance in natural T-IELs. Thus, the strong and sustained TCR signals that T-IEL precursors receive during agonist selection drive a unique developmental trajectory that equips them with the unconventional signaling properties required for their specialized role in the gut.

### LAT downregulation is a key component of RePrESS

Our phosphoproteomic analysis revealed that natural T-IELs exhibit unique signaling characteristics, particularly an enrichment of phosphopeptides from several negative regulators of TCR signaling, including receptors CD200R1 and 2B4/CD244 ^35^, adaptors DOK1 ^36^ and LAT2 ^37^, and tyrosine phosphatases SHP-1 (encoded by *Ptpn6*) and SHP-2 (encoded by *Pptn11*) compared to conventional CD8 T cells **(Fig. 3f and Suppl. Fig. 5)**. SHP-1 and SHP-2 can dephosphorylate critical TCR signaling intermediates, such as CD3ζ and ZAP70 ^38^, and are highly expressed in natural T-IELs **(Suppl. Fig. 7a)**. However, deletion of both SHP-1 and SHP-2 in T-IELs did not alter ERK or S6 phosphorylation upon *in vitro* anti-CD3 stimulation nor IFNγ production after *in vivo* TCR stimulation **(Suppl. Fig. 7b-d)**. LAT2 has been suggested to suppress TCR signaling by sequestering key signaling molecules away from LAT ^37,39,40^, and was previously implicated as a potential explanation of natural αβ T-IEL hyporesponsiveness ^18^. Using T-IELs from *Lat2* knockout (*Lat2^-/-^*) mice, we evaluated whether the absence of LAT2 could alleviate the suppressed TCR signaling observed in natural T-IELs. There was no effect of loss of LAT2 on T-IEL numbers **(Suppl. Fig 8a)**, and on ERK and S6 phosphorylation after in vitro stimulation with anti-CD3 **(Suppl. Fig. 8b)**. However, *in vivo* stimulation with anti-CD3 elicited significant increases in production of cytokines IFNγ and TNF in the natural T-IEL subset. These effects were not observed in splenic T cells, where LAT2 is not expressed **(Suppl. Fig. 8c-d)**. Despite these findings, the modest magnitude of the cytokine response suggested that LAT2 alone is not the primary driver of TCR hyporesponsiveness in natural T-IELs. Thus, these molecules may be only part of a suite of inhibitory mechanisms that ensure T-IEL tolerance.

LAT is critical for assembling the signalosome downstream of TCR engagement, and its absence or reduced activity is known to impair TCR signaling ^41,42^. As LAT is downregulated in natural T-IELs, we hypothesized that restoring LAT expression could rescue TCR signaling and validate its role as a central component of the rewiring of natural T-IELs. To test this, we retrovirally transduced T-IELs with a LAT-IRES-GFP construct or GFP control. After 7 days in culture, LAT expression was restored in ∼15% of T-IELs **(Fig. 5a)**. Upon anti-CD3 stimulation, we observed significant increases in the phosphorylation of key signaling molecules, including CREB, ERK, and S6, in natural T-IELs transduced with LAT compared to GFP controls **(Fig. 5b-c)**. Notably, this effect was restricted to natural T-IELs, as induced T-IELs did not exhibit similar improvements in TCR signaling upon LAT expression **(Fig. 5b-c)**. Importantly, the rescued signaling with LAT expression also correlated with increased functional responses, as a significantly larger proportion (up to 50%) of GFP-LAT transduced natural T-IELs produced IFNγ compared to GFP-transduced cells, indicating tolerance was reversed **(Fig. 5d)**. Overall, our findings demonstrate that tolerance in natural T-IELs is driven by a distinct mechanism of hyporesponsiveness to TCR stimulation that is elicited through global rewiring of the TCR signalosome involving both LAT downregulation and LAT2 expression.

**Figure 5.**
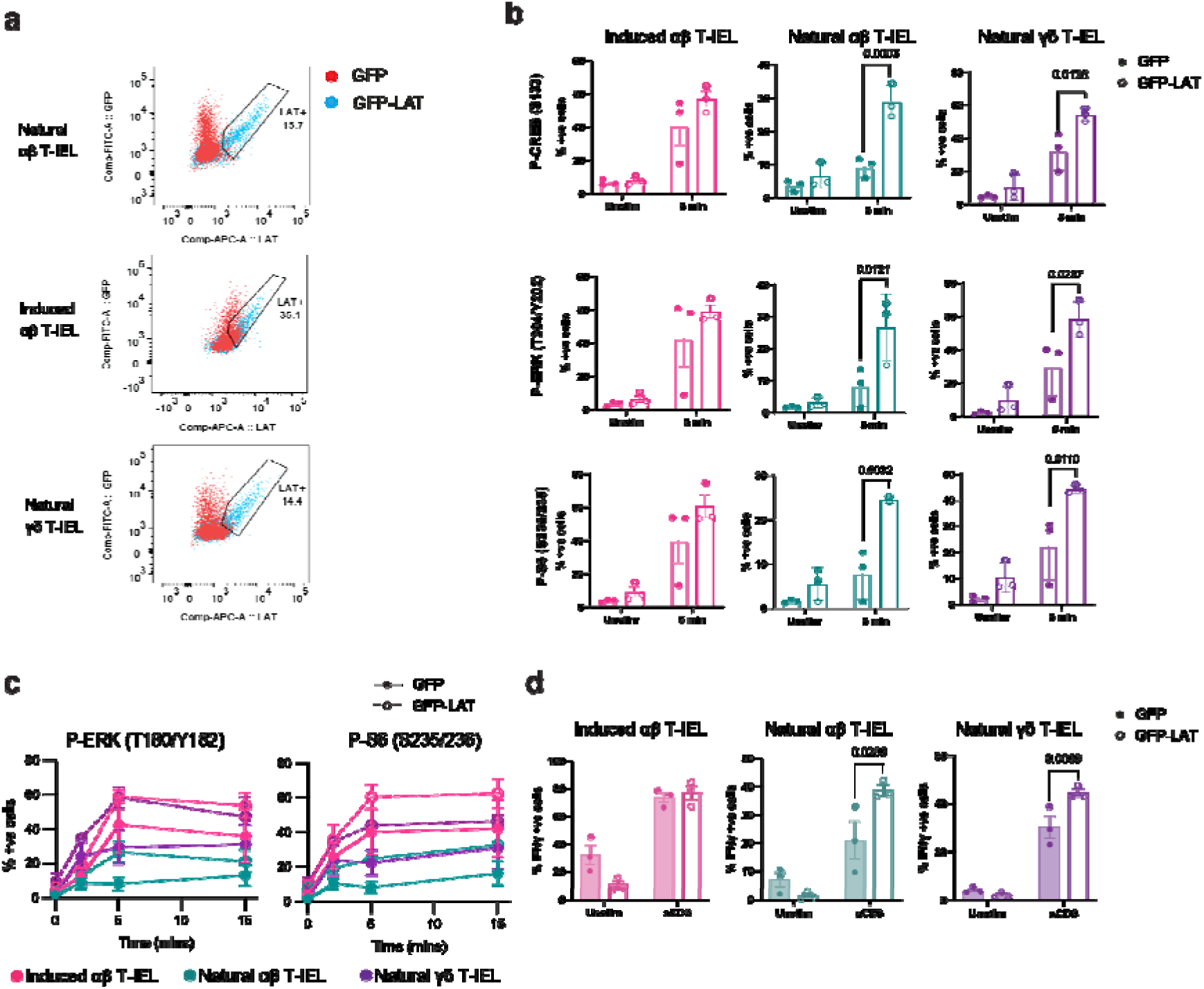
Forced expression of LAT restores TCR functionality in natural T-IEL. (a) Representative flow cytometry dot plots showing GFP and LAT expression in T-IEL subsets transduced with either GFP empty vector (red) or LAT-IRES-GFP (blue). Gated regions indicate the percentage of LAT-positive cells. (b) Bar graph depicting the percentage of natural T-IEL subsets positive for phosphorylated CREB, ERK, and S6 following 5 minutes of anti-CD3 stimulation in cells transduced with either GFP empty vector (filled circles) or LAT-IRES-GFP (empty circles). Data represent mean ± SEM from three biological replicates. (c) Time-course analysis of ERK and S6 phosphorylation following anti-CD3 stimulation in T-IEL subsets transduced with either GFP empty vector (solid lines, filled circles) or LAT-IRES-GFP (dotted lines, empty circles). (d) Percentage of T-IEL subsets positive for intracellular IFNγ following anti-CD3 stimulation, comparing cells transduced with GFP empty vector (filled circles) vs. LAT-IRES-GFP (empty circles). Data represent mean ± SEM from three biological replicates. Statistical significance in (b) and (d) was determined using ANOVA with Tukey’s post hoc correction.

### RePrESS is a conserved mechanism broadly underpinning innate-like T cell tolerance

The high level of congruency of natural T-IEL signalosome signature (RePrESS) between phosphoproteomic, proteomic and transcriptomic data of mouse T-IELs provided an opportunity to leverage publicly available single cell transcriptomic datasets to evaluate the conservation of this TCR signalosome signature in other species. Our analyses of human ^33^ and porcine natural T-IELs ^43^ revealed that this signature is evolutionarily conserved in humans and pigs, highlighting its fundamental role in T-IEL biology **(Fig. 6a,b)**. Notably, several of the RePrESS gene signature were also shown to define the transcriptional profile of human agonist-selected CD8αα^+^ T-IEL precursors in the thymus, i.e., *GCSAM, PLCG2, CLNK*, among others ^44^, indicating human T-IELs are also rewired during thymic maturation.

**Figure 6.**
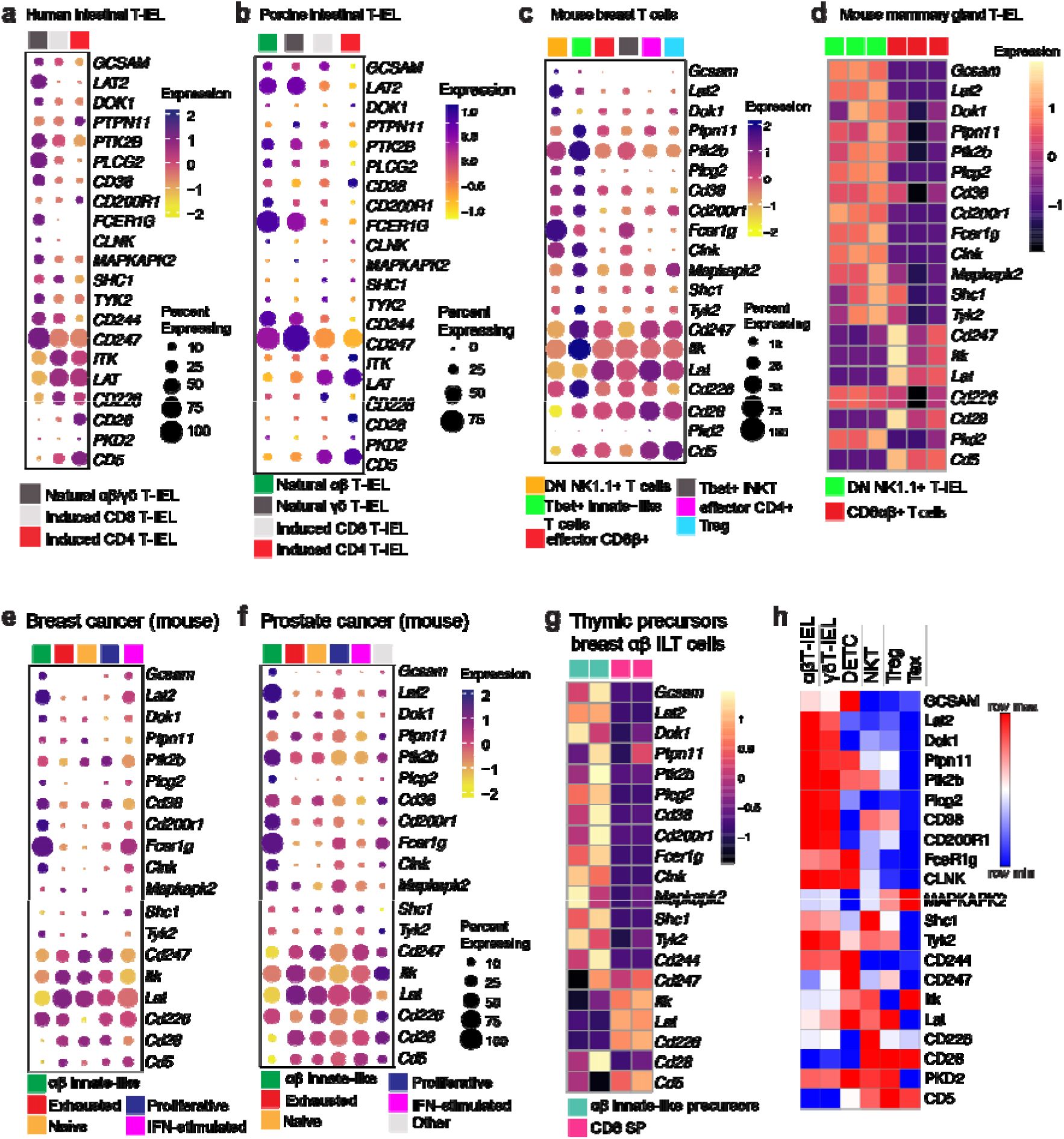
The RePrESS gene signature is evolutionarily conserved and broadly applicable to innate-like T cells. Bubble plots showing gene expression levels and frequency of cells expressing natural T-IEL signalosome RePrESS signature genes in (a) human^33^ and (b) porcine^43^ natural T-IEL clusters, from published single-cell RNA sequencing (scRNA-seq) datasets. (c) Bubble plots showing gene expression levels and frequency of cells expressing the RePrESS signature genes in T-bet+ T cells isolated from breast tissue of nulliparous, pregnant, and lactating mice. (d) Heatmap of the RePrESS signature genes in bulk RNA seq analyses of mammary gland natural CD4-CD8β-(DN) CD8α+ NK1.1+ T-IELs, compared with mammary CD8αβ+ T-IEL. (e-f) Bubble plot analysis of the RePrESS gene signature in scRNA-seq data from (e) murine breast cancer (PyMT mice) and murine prostate cancer (TRAMP mice) tissues^46^. (g) Heatmap of the RePrESS gene signature in thymic conventional CD8 single positive (SP) progenitors and αβ innate-like T (ILT) cell progenitors. (h) Row-normalized heatmap of natural T-IEL TCR signalosome signature protein intensities in natural αβ and γδ T-IEL, dendritic epidermal T cells (DETC), liver invariant natural killer T (iNKT) cells, natural regulatory T cells (nTreg) T and exhausted T cells (Tex), extracted from quantitative total proteomics measurements.

During pregnancy and lactation, an increase in T-bet-expressing T cells were found within the mammary glands of female WT mice ^45^. Analysis of published scRNA-seq of these T cells revealed two innate-like T-IEL populations, DN NK1.1+ and T-bet+ innate-like T cells, both of which were enriched in the RePrESS gene signature **(Fig. 6c)**. Further analyses of bulk transcriptomics data of the DN NK1.1+ natural T-IEL compared to conventional CD8β+ T cells from mammary glands of pregnant mice, confirmed that this mammary innate-like T-IEL population express the identical RePreSS gene signature as intestinal natural T-IEL **(Fig. 6c)**. Similar innate-like T cells were also found in breast tumor tissue from PyMT mice (mouse mammary tumour virus-PyMT) and prostate cancer tissue from TRAMP mice ^46^. In these innate-like T cells too, the RePreSS signature was highly conserved **(Fig. 6d-e)**. Interestingly, exhausted T cells from these tumors did not share the same gene signature, supporting the notion that chronic TCR stimulation driving exhaustion does not induce TCR hyporesponsiveness through the same TCR signalosome rewiring. We also reanalyzed bulk RNA-seq data from the thymic precursors of the innate-like αβT-IEL found in breast cancer ^46^ and found that the RePreSS gene signature was already differentially expressed in these post-selection thymic progenitors of breast innate-like αβT cells, when compared to CD8 SP (**Fig. 6f**).

Since agonist selection induces the RePrESS signature in natural αβΤ-IEL, we analyzed the proteomes of other agonist-selected T cells, including liver NKT cells and splenic natural Tregs, as well as other innate-like T cells such as skin dendritic epidermal T cells (DETC) for the RePrESS proteins. While DETC share upregulation of certain proteins such as GCSAM, LAT2, PLCγ2, FcεR1γ and CLNK, LAT was not downregulated **(Fig. 6g)**. Similar findings were seen in iNKT and nTregs. We also analyzed the proteomes of exhausted T cells (Tex) generated by repeated TCR restimulation ^47^. Tex did not show the upregulation of the TCR proteins specific to the natural T-IEL signalosome, and expressed no or very low levels of most of the B cell/myeloid cell proteins seen in natural T-IELs, as in the tumor tissue. However, exhausted T cells did show a strong downregulation of LAT, which may be critical for their lack of TCR responses. Altogether, the distinct mechanism of tolerance we have identified in intestinal natural T-IEL is highly conserved in evolution, and broadly applicable to other self-reactive T cells, particularly innate-like T cells in non-lymphoid and barrier tissues.

### Loss of the RePrESS gene signature in autoimmune T-IEL in Celiac disease

As the RePrESS gene signature is highly conserved in human natural intestinal T-IEL, we hypothesized that changes in this rewiring might contribute to autoimmunity. In celiac disease (CD), recognition of gluten peptides from the diet by lamina propria CD4+ T cells in susceptible individuals leads to an abnormal immune response characterized by massive destruction of the intestinal epithelium^48^. While the CD4 T cells are the initiators of the disease, it is the massive expansion of cytotoxic T-IELs that drive the enterocyte destruction. It was recently shown that the major expansion occurs in the natural T-IEL populations in active disease, and these expanded natural T-IEL take on a more cytotoxic effector-like profile, implicating them as drivers of the autoimmune damage^6^. This expanded cytotoxic population are referred to as natural-effector T-IEL (T_NE_) and are largely absent in healthy controls, but share many similar TCR clonotypes with the healthy natural T-IEL populations. We therefore analyzed previously published scRNA seq data of CD8+ T-IEL from healthy controls and patients with active (ACD), potential (PCD), or on a gluten-free diet (GFD), to assess changes in the RePrESS gene signature in natural healthy and effector T-IEL populations. In this dataset as well, the natural αβT-IEL population clearly express the RePrESS signature (**Fig. 7a**). However, in ACD, the expression pattern of the whole gene set is changed in the natural T-IEL. Further, in the effector natural T-IEL population found mainly in ACD, it is even clearer that the RePrESS gene signature is not expressed. Thus, under conditions of autoimmunity, the TCR signalosome is no longer repressed, allowing autoimmune activation of natural T-IEL.

**Figure 7.**
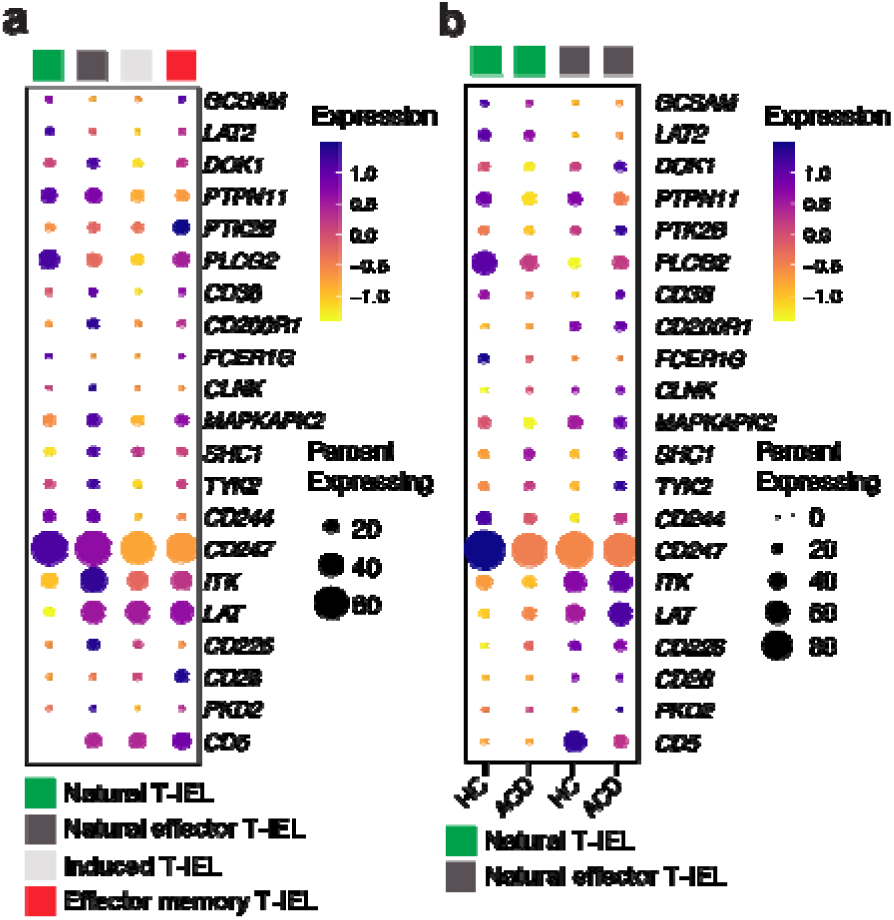
The RePrESS gene signature is lost in human celiac disease natural intestinal T-IEL. Analyses of published^6^ scRNA-seq data of human duodenal CD8 T-IEL from healthy controls (HC, n=17), and patients with potential celiac disease (PCD, n=7), active celiac disease (ACD, n=11), or celiac disease on a gluten-free diet (GFD, n=19). (a) Bubble plots showing gene expression levels and frequency of cells expressing the TCR signalosome RePrESS signature genes in natural, natural effector, induced, and induced effector CD8+ αβT-IEL populuations from pooled data from all conditions. **(b)** Bubble plots showing gene expression levels and frequency of cells expressing the RePrESS signature genes in natural αβT-IEL, and natural effector αβT-IEL, comparing cells from HC vs patients with ACD. Note that the natural effector αβT-IEL population represents <5% of total CD8+ T-IEL in HC, and represents 10-50% of total CD8 T-IEL in ACD.

## Discussion

The persistence of self-reactive T cells in peripheral tissues necessitates robust mechanisms of immune tolerance to prevent autoimmunity ^54^. In this study, we demonstrate that self-reactive natural T-IELs are not controlled by traditional tolerance strategies such as Treg-mediated immunosuppression, anergy, or exhaustion. Instead, we uncover a distinct mechanism of tolerance that involves developmental changes in the expression of TCR signaling proteins, which we term RePrESS (Rewiring of Proximal Elements of the TCR Signalosome for Suppression). This mechanism enables autoreactive natural αβ T-IELs to tolerate self-ligands by eliciting TCR hyporesponsiveness through rewiring of their proximal TCR signaling machinery. Thus, we show that RePrESS-mediated tolerance in natural T-IELs is an evolutionarily conserved and programmed developmental process, that is tightly regulated and breaks down in autoimmune diseases such as celiac disease.

Why do natural T-IELs require this specialized mechanism? Unlike circulating T cells, T-IELs reside in constant, intimate contact with epithelial cells that express cognate self-ligands, such as non-classical MHC molecules for intestinal αβ T-IELs. In this context, traditional Treg-mediated tolerance may be insufficient, especially considering the scarcity of Tregs in the intestinal epithelium. Other tolerance mechanisms such as anergy or exhaustion would impair T-IEL functionality altogether. Instead, RePrESS offers a developmentally hardwired mechanism of restraint that allows T-IELs to physically interact with their ligands while remaining non-inflammatory. This is likely a critical adaptation to their immunosurveillance function - natural T-IELs must constantly patrol the epithelial barrier, detect stress, and respond to pathogens—yet without being triggered by the abundant self and commensal-derived antigens in their niche. By decoupling TCR engagement from full activation, RePrESS allows natural T-IELs to remain vigilant but silent until appropriately activated through innate cues, such as stress receptors or cytokine receptors.

Specifically, we demonstrate that RePrESS involves downregulation of LAT and CD28, alongside the upregulation of alternative molecules such as NTAL/LAT2, FcεR1γ, Lyn, and CLNK (**Suppl. Fig. 9**). This remodeling emerges during thymic development following agonist selection, where strong and sustained TCR signals appear to drive reprogramming. Notably, lack of CD28 co-stimulation favors development of natural T-IELs ^55^. Potentially, very strong TCR signals in the absence of costimulation leads to the downregulation of key TCR signaling molecules, and to the concomitant expression of several non-T cell proteins, including LAT2, Lyn, CLNK, and GCSAM, that are normally expressed in B cells. Interestingly, forced expression of LAT in bone marrow progenitors blocks B cell development^56^. Hence, it is possible that LAT expression represses B cell lineage proteins, whereas in natural T-IELs, reduction of LAT permits their re-expression. As RePrESS involves a whole program of rewiring, it suggests the involvement of a master transcriptional regulator. Potential candidates include Id2, Id3 and Ikzf2 that are highly expressed in agonist-selected T-IEL precursors^34^. However, deciphering the contributions of these factors is confounded by the ablation of T-IEL in mice lacking Id2 and Id3^34,57^. Importantly, this programmed rewiring in natural T-IELs occurs during thymic selection rather than clonal escape, defining this process as a new mechanism of non-deletional tolerance.

Surprisingly, canonical inhibitory regulators of TCR signaling such as SHP-1/2 and LAT2 are largely dispensable for maintaining TCR hyporesponsiveness in T-IELs. Instead, LAT downregulation partially explains the lack of TCR responses, underscoring the requirement for a coordinated global remodeling of the TCR signalosome. The reliance on LAT downregulation may be unique to innate-like T cells, as loss LAT impairs the survival and function of agonist-selected Tregs^41^. Interestingly, in active celiac disease, LAT is highly upregulated in natural T-IEL, suggesting LAT may be potentiating autoimmune responses in this context. The autoimmune upregulation of LAT in celiac disease may be driven by chronic signaling through stress receptors such as NKG2D and CD94, as well as IL-15 that is highly expressed in active disease^48^. Conversely, the loss of LAT in exhausted T cells may be important for their lack of responsiveness in cancer immunotherapy. Understanding the mechanisms of TCR signaling rewiring may thus be important not only for regulating autoimmunity, but also for rescuing TCR responses in cancer.

Does T-IEL hyporesponsiveness imply that natural T-IEL do not need their TCR after thymic selection? Indeed, ablation of the TCR on natural αβΤ-IEL does not affect their survival or maintenance of their phenotypes^59^. Yet a mouse model of intestinal inflammation, the self-specific TCR of natural αβT-IEL and cognate ligand expression in the intestine was required for their ability to prevent colitis^8^. Similarly, Btnl1-selected Vγ7+ natural T-IEL need expression of Btnl1 on the intestinal epithelium for selection, but not for long-term maintenance^60^. However, in the absence of the TCR binding Btnl1 molecule, CD122 was downregulated on Vγ7+ natural T-IEL, implying there is still a functional role of the TCR on these cells. DETC do not need TCR signaling through LAT for their long-term survival and maintenance, but need LAT for their response to skin wounding^58^. DETC have previously been shown to have a very high TCR activation threshold^20^, yet TCR signaling is required for the expression of innate stimulatory receptors on DETC, allowing DETC to respond rapidly to epithelial stress^53^. These findings lead to the proposition that DETC TCR signaling is important for normality sensing. These seemingly contradictory results may be reconciled if we consider rewiring of TCR signaling couples TCR binding to alternate functions. Thus, rewiring prevents IFNγ production and cytotoxicity, but allow expression of innate inhibitory and stimulatory receptors, and production of immunoregulatory molecules such as IL-10.

These insights raise the exciting possibility that RePrESS represents not just a mechanism of natural tolerance, but a targetable axis for therapeutic modulation. Understanding how proximal TCR signaling is rewired opens up new avenues to either restore TCR responsiveness in settings of dysfunction (e.g., cancer, chronic infection), or to induce a RePrESS-like state in autoreactive T cells in autoimmune diseases. For instance, therapeutic modulation of LAT expression or downstream signaling adaptors could potentially recalibrate self-reactive T cells without inducing global immunosuppression. Furthermore, the RePrESS program offers a new framework to interpret T cell behavior in tissues beyond classical paradigms of anergy, exhaustion, or suppression. This could inform the design of tissue-adapted T cell therapies, including engineered TCR or CAR-T cells intended to operate in epithelial or tumor environments, where inappropriate activation risks off-target toxicity.

Our findings suggest that RePrESS represents a developmentally programmed mechanism of tolerance induction in natural T-IELs. While we focused primarily on ‘type a’ agonist-selected natural αβ T-IELs ^61^, this mechanism likely extends to other innate-like lymphocyte subsets, including γδ T-IELs and DETCs, as we show here. The rewiring of TCR signaling may support their immunosurveillance behavior ^53,62^ and further allow these T cell responses to be driven by other activating receptors and cytokine receptors. Thus, our study provides a conceptual framework for understanding how T-IELs and other self-reactive T cells can persist in the periphery without triggering autoimmunity.

## Limitations of the study

While this study has revealed remodeling of the TCR signalosome as a mechanism for dampening conventional TCR signaling in innate-like lymphocytes, we have not addressed whether this rewiring drives an alternate functional response. We have shown in vitro that restoration of LAT expression rescues functional responses, but whether this would lead to autoreactivity in vivo is not clear. Further studies will also be needed to identify the transcriptional drivers of the RePrESS program, to allow modulation of these changes.

## Acknowledgements

We are very grateful to Dr. I. Moraga (University of Dundee) for help with fluorescent cell barcoding set up and Profs. D. Cantrell and Y. Kulathu (University of Dundee) and Prof. S. Minguet (University of Freiburg) for many useful discussions. We thank Dr. Russell S. Hamilton, AstraZeneca, Cambridge, for guidance and advice on scRNAseq analyses. We acknowledge the contribution of Dr. V Bugajev and Prof. Petr Draber (Prague, Czech Republic) for initial analyses of the LAT2 KO mice. We thank the NIH Tetramer Facility for providing CD1d tetramers. We would like to thank A. Rennie and R. Clarke for cell sorting, the staff of the Biological Resource Unit for technical assistance, and the mass spectrometry facility at the MRC Protein Phosphorylation and Ubiquitylation Unit. We also acknowledge help of Dr. Gemma Alderton, Biosciedit, for critical reading and editing of the manuscript.

## Author contributions

MS conceptualized and designed the study with input from ASC. HJW, ASC, SAS, NS, IS, KW, MO, JS, FL, SP, EK, KDR, AC, DB, MS performed experiments and analyzed data. CP, PGT, KK, DB and MS provided funding and conceptual and supervisory inputs. ASC, HJW, and MS prepared figures and wrote the manuscript with input from all co-authors.

## Methods

### Ethics

Mice were bred and maintained with approval by the University of Dundee ethical review committee under a UK Home Office project license (PD4D8EFEF, PP2719506) in compliance with UK Home Office Animals (Scientific Procedures) Act 1986 guidelines. Nur77-Tempo mice were bred and maintained at the University of Birmingham under UK Home Office Project Licence P18A92E0A. Foxp3-DTR mice were bred and maintained in SPF conditions at Cincinnati Children’s Hospital Medical Center by protocols approved by IACUC. *foxp3a*^-/-^ and *foxp3a*^+/+^ zebrafish husbandry and experiments were performed in accordance with National Cerebral and Cardiovascular Center Research Institute, Japan ethics and national animal guidelines.

### Mice

C57BL/6J mice were purchased from Charles River, UK and acclimatized for at least 7 days prior to use. Ly5.1 mice were purchased from Charles River and bred in house. *Lat2^-/-^* (LAT2 KO) mice were obtained from Prof. Petr Draber (Prague, Czech Republic) under MTA from Prof. Bernard Malissen (Marseilles, France). *Ptpn6^fl/fl^* (JAX stock #008336) or *Ptpn6^fl/fl^/Ptpn11^fl/flBgn^* mice ^52^, B6.129(Cg)-*Foxp3^tm4(YFP/icre)Ayr^*/J (JAX stock 016959) were obtained from Doreen Cantrell, University of Dundee and crossed to *GzmB^Cre/+^* (Gzmb-Cre knockin) mice that were generated by Ingenious Targeting Laboratory for MS. This resulted in generation of GzmB-Cre/ *Ptpn6^fl/fl^* and GzmB-Cre/ *Ptpn6^fl/fl^/Ptpn11^fl/fl^* mice that had deletion of SHP-1 (*Ptpn6*) alone or SHP-1 and SHP-2 (*Ptpn6* and *Ptpn11*) in all Granzyme B expressing cells. As T-IEL highly express Granzyme B, *Ptpn6* and/or *Ptpn11* were deleted in these cells. Mice were maintained in a standard barrier facility on a 12hr light/dark cycle at 21°C in individually ventilated cages with sizzler-nest material and fed an R&M3 diet (Special Diet Services, UK) and filtered water ad libitum. Cages were changed at least every 2 weeks. Nur77-Tempo mice were bred and maintained at the University of Birmingham. Foxp3-DTR mice^24^ (JAX stock #016958) were bred and maintained in SPF conditions at Cincinnati Children’s Hospital Medical Center by protocols approved by IACUC. For Treg depletion experiments, Foxp3-DTR mice were given diphtheria toxin (DT) 25μg/kg on day 0 and 10μg/kg subsequently on day 3 (diluted in PBS) intraperitoneally. Age- and sex-matched mice between 6 and 12 weeks were used for all experiments. Mice of both genders were used. No statistical methods were used to pre-determine sample size, but our sample sizes are similar to those reported in previous publications.

### Zebrafish analyses

*foxp3a*^-/-^ and *foxp3a*^+/+^ zebrafish husbandry and experiments were performed in accordance with National Cerebral and Cardiovascular Center Research Institute, Japan ethics and national animal guidelines. Animal density was maintained at 3 to 5 fish L−1. Clutch mates were used as controls in all experiments. For Quantitative reverse transcriptase polymerase chain reaction (RT-qPCR), total RNA from fish organs was extracted using TRIzol reagent. cDNA was subsequently synthesized using a SensiFast cDNA synthesis kit (Meridian). PCR was performed using a CFX Connect realtime PCR system (Biorad). The amount of cDNA was normalised to *actb2*/β*-actin2* expression in RT-qPCR experiments. The primers used in this study are listed in supplementary table 3.

### Isolation of Intestinal Intraepithelial T Lymphocytes

T-IEL were isolated as previously described ^63^. Briefly, the small intestine was removed and flushed with PBS. The intestine was cut longitudinally and then transversely into 0.5 cm pieces. Gut pieces were incubated in RPMI containing 10% FBS, Penicillin/Streptomycin, L-Glutamine and 1 mM DTT. After 40-minute incubation with shaking, pieces of gut were vortexed and filtered through a 100 μm cell strainer. Filtrate was spun and resuspended in 44% Percoll. This was layered on top of 75% Percoll (Sigma) and spun at 700g for 30 minutes without brake. Cells were removed from the interface of the Percoll layers and washed before further use.

### Isolation of thymocytes, splenocytes and lymph node cells

Thymus, spleen or pooled inguinal, mesenteric, brachial, axillary and cervical lymph nodes were removed from mice and crushed through a 70 μm cell strainer into isolation media. Cell suspensions resuspended in appropriate buffer for downstream applications.

### Stimulation and Fluorescent Cell Barcoding with Amine reactive dyes

T-IEL were isolated and sorted with the CD8a Positive Selection Kit (Stemcell Technologies) and stained with Live/Dead Near-Infrared Amine reactive dye (ThermoFisher Scientific) then resuspended in stimulation media at 1 x 10^6^ cells/ml. Lymph node CD8α^+^ T cells were isolated and sorted using the CD8a Positive Selection Kit (Stemcell Technologies). Isolated cells were washed and resuspended at 1 x 10^6^ cells/ml in stimulation media. Stained T-IEL and unstained lymph node T cells were mixed at a 1:1 ratio before stimulation with 30 μg/ml anti-CD3 (Clone 145-2C11, Biolegend) crosslinked with 5 μg/ml anti-Hamster (Jackson Immunoresearch). PP2 was added at 20 μM, GDC-0941 at 1 µM and Rapamycin at 20 nM to relevant condition for 1 hour prior to stimulation, pervanadate was used at 10 μM. PMA was used at 100 ng/ml and ionomycin at 1 μg/ml. During stimulation cells were incubated at 37°C on shaking incubator, 400 rpm. Cells were fixed in 2% paraformaldehyde (PFA), incubated for 10 minutes, and permeabilized with 90% ice cold methanol.

After permeabilization, the different stimulation conditions were subject to barcoding with different concentrations of Pacific Blue NHS ester dye or Pacific Orange succinimidyl ester (both ThermoFisher Scientific). Barcoded conditions were pooled and then subject to further intracellular and surface staining.

### Phospho-flow cytometry

Fixed and permeabilized cells were incubated with primary phospho-antibodies; Phospho-CD3ζ (Y142) (Clone K25-407.69), Phospho-SLP-76 (Y128) (Clone J141-668.36.58) (both BD Biosciences); Phospho-ZAP-70 (Y319)/Syk (Y352) (Clone 65 E4), Phospho-ERK1/2 (T202/Y204) (Clone 197G2), Phospho-Akt (S473) (Clone 193H12), Phospho-S6 ribosomal protein (S235/236) (Clone D57.2.2E), Phospho-4EBP1 (T37/46) (Clone 236B4), Phospho-CREB (S133) (Clone 87G3), Phospho-p38 MAPK (T180/Y182) (Clone D3F9), P-Akt (T308) (Clone D25E6), Phospho-p70 S6 kinase (T389), Phospho-PI3K p85 (Y458)/p55(Y199) (all Cell Signaling Technology). Primary antibodies were incubated for 30 minutes at room temperature in the dark before incubation with secondary antibody, anti-rabbit Dylight649 (Biolegend) or Alexa Fluor 647 AffiniPure (Fab’)_2_ Fragment Donkey Anti-Mouse IgG (H+L) (Jackson ImmunoResearch). Secondary antibody was incubated for 30 minutes at room temperature in the dark before washing and incubation with surface stains; TCRβ-PE (Clone H57-587) and CD8α-PE-Cy7 (Clone 53-6.7) (both Biolegend); TCRγδ-PerCP efluor710 (Clone GL3) and CD8β-FITC (Clone H35-17.2) (both eBioscience). After incubation, cells were washed and read on Cytoflex (Beckman Coulter).

### Flow cytometry

Intracellular staining was performed on cells isolated as described above that were stained with Live/Dead Fixable Near-InfraRed or Live/Dead Fixable Blue (1:1000) (ThermoFisher Scientific) and fixed and permeabilized using the Fixation/Permeabilisation and Permeabilization buffers (eBiosciences) following manufacturers protocols. Permeabilized cells were stained with antibodies, LAT (1:200), SHP-1 or SHP-2 (all Cell Signaling Technology) followed by anti-rabbit Dylight649 (1:500) (Biolegend), NTAL-PE (1:100) (ThermoFisher Scientific).

Surface staining was performed on cells. Antibodies used were CD1d-APC (1:200), CD4-BV605 (1:400), CD8a-AF700 (1;400), CD5-PE-Cy7 (1:400), CD25-APC (1:100), CD122-PE (1:200), PD-1-FITC (1:200) (all Biolegend) and TCRb-BV786 (1:200) (BD Biosciences).

T-IEL, spleen/LN cells, thymocytes isolated from untreated Nur77-Tempo or WT mice that were injected with anti-CD3 (Clone 145-2C11) or IgG (both Biolegend) were stained with Viability dye-efluor780 (eBiosciences), then TCRβ-AF700 (Clone H57-597), TCRγδ-APC (Clone CL3), CD8β-PerCP-Cy5.5 (Clone 53-5.8) (all Biolegend); CD8α-BUV395 (Clone S3-6.7), PD-1–PE-Cy7(Clone A1718BB) and CD4-BUV737 (Clone GK1.5) (both BD Biosciences). Fluorescence of Nur77-blue and Nur77-red in all T cell subsets was ascertained by flow cytometry.

T-IEL and splenocytes isolated from untreated Foxp3-DTR mice injected either with PBS or DT were stained with Viability dye Zombie Yellow (Invitrogen), CD45 BV785 (clone 30-F11), CD103 BV421 (Clone 2E7), TCRβ AF700 (Clone H57-597), TCRγδ BV510 (Clone CL3), CD4 BV711 (Clone GK1.5), CD8α APC Fire (Clone S3-6.7), CD8β PerCP (Clone 53-5.8), GzmA PE (Clone 3G8.5), CD44 BV650 (Clone IM7), IFNγ APC (Clone XMG1.2), TNF PE (Clone MP6-XT22). All the antibodies were from Biolegend.

### Cytokine production upon *in vivo* stimulation with anti-CD3

Mice were treated with 50 µg anti-CD3 Ultra-LEAF (Clone 145-2C11) (Biolegend) or sterile PBS (Gibco) for 3 hours prior to harvesting T-IEL and spleen. Tissues were processed following protocols described above. Cells were treated with Brefeldin A and GolgiStop (both Biolegend) for 2 hours prior to fixation and permeabilization with 1X Fixation/Permeabilisation buffer and 1X permeabilization buffer (eBiosciences). Cells were stained with TNFa-PE (1:200), IFNg-PE Dazzle594 (1:200), CD8a-PE-Cy7 (1:400), CD8b-FITC (1:400) and CD4-BV605 (1:400) (all Biolegend), TCRb-BV786 (1:200), TCRgd-PerCP-ef710 (1:200) (both BD Biosciences). Cells were acquired on LSR Fortessa.

### Zebrafish Histology and Imaging

Whole zebrafish were euthanized and fixed in Davidson’s Fixative (Muto Pure Chemicals, Japan) at 4°C overnight. Fixed specimens were embedded in paraffin, and 10-μm sections were obtained for hematoxylin-eosin (H&E) staining, following standard protocols (Applied Medical Research, Japan). H&E-stained sections were visualized using a Nanozoomer S60 microscope (Hamamatsu, Japan).

### Foxp3-DTR Mice Histology and Imaging

Foxp3-DTR mice were euthanized, followed by systemic perfusion with PBS to remove circulating blood. Tissues were then excised and fixed in 4% PFA at 4°C overnight. Fixed tissues were processed for paraffin embedding, and 5-μm sections were prepared and stained with H&E using standard protocols. Imaging was performed on a Nikon Eclipse Ni-U microscope.

### T-IEL Cultures

T-IEL were isolated and purified using the CD8α Positive Selection kit (StemCell Technologies) following manufacturer’s instructions. Cells were resuspended in RPMI(Gibco) supplemented with 10% FBS, penicillin, streptomycin, 2 mM L-Glutamine, 1 mM sodium pyruvate, 1 mM β-mercaptoethanol, 2.5 mM HEPES and 1% non-essential amino acids. Recombinant murine IL-15 (Peprotech) was added at 20ng/ml.

### Degranulation assay

T-IEL were isolated and purified using the CD8α Positive Selection kit (StemCell Technologies) following manufacturer’s instructions. Cells were cultured as indicated before plating at 1x10^6^ cells/ml in culture media supplemented with Brefeldin A (Biolegend), BD GolgiStop (BD Biosciences), CD107a (1D4B) and CD107b (M3/84) (both Biolegend). Cells were plated in a 96-well round-bottom plate coated with 3 μg/ml anti-CD3 (Biolegend), uncoated well or with PMA (1 ng/ml) and Ionomycin (1 μg/ml). Cells were harvested after 4 hours and fized with Fixation/Permeabilization Concentrate and permeabilized with Permeabilisation buffer (eBiosciences). Cells were then stained for IFNγ-PE/Dazzle 594 (CloneXMG1.2) (Biolegend) and fluorescence measured on LSR Fortessa (BD Biosciences).

### Sample preparation for Phospho-proteomics

T-IEL and lymph node cells were isolated following the previously described method. CD4 and CD8b cells were depleted from the T-IEL compartment using CD4 and CD8b biotinylated antibodies and magnetic streptavidin nanobeads (MojoSort) (Biolegend). CD8a^+^ cells were purified from CD4 and CD8b depleted T-IEL and lymph node T cells using the CD8a Positive Selection Kit (Stemcell Technologies). Cells were resuspended in RPMI + 1% FBS + L-Glutamine + Pen/Strep at a concentration of 1x10^6^ cells/ml and stimulated with 10 µM of pervanadate. After 5 minutes cells were pelleted and snap frozen. Cells were lysed with 5% sodium dodecyl sulphate + 50 mM triethylammonium bicarbonate (TEAB) (Sigma). Samples were lysed by rounds of shaking at 1000 rpm at room temperature and at 95°C at 500 rpm and sonication. Benzonase was added and incubated for 15 minutes at 37°C. Samples were prepared following the S-trap mini protocol (Protifi). Briefly, lysates were reduced using 10 mM TCEP, alkylated with 0.5 M chloracetamide, acidified with 12% phosphoric acid. 300 µg of each sample was loaded onto an S-Trap mini column and digested using trypsin (Sigma) at 1:20 wt:wt. Samples were eluted from the column and vacuum dried using SpeedVac. Samples were subject to C18 Clean up (Waters) and labelling with 10-plex Tandem Mass Tagging Kit (ThermoFisher Scientific) following manufacturers protocols. Labelled samples were combined and vacuum dried. Samples were resuspended in 50 mM ammonium bicarbonate and incubated with SH2 superbinder beads (Precision Proteomics). After 1 hour of incubation beads were washed thoroughly with ammonium bicarbonate. Beads were resuspended in 0.4% trifluoroacetic acid (TFA) then passed through a Spin-X column (Corning). Samples were vacuum dried before resuspension in formic acid.

The quenched samples were then mixed and fractionated with high pH reverse-phase C18 chromatography using the Ultimate 3000 high-pressure liquid chromatography system (Thermo Scientific) at a flow rate of 500 µl/min using two buffers: buffer A (10 mM ammonium formate, pH 10) and buffer B (80% ACN, 10 mM ammonium formate, pH 10). Briefly, the TMT-labelled samples were resuspended in 200 µl of buffer A and desalted then fractionated on a C18 reverse-phase column (4.6L×L250Lmm, 3.5 µm, Waters) with a gradient as follows: 3% Buffer B for 19 min at 275 µl/min (desalting phase), ramping from 275 µl/min to 500 µl/min in 1 min, 3% to 12% buffer B in 1 min, 12% to 40% buffer B in 30 min, 40% B to 60% B in 5 min, 60% B to 95% B in 2 min, 95% for 3 min, ramping to 3% B in 1 min and then 3% for 9 min. A total of 96 fractions were collected and then concatenated into 12 fractions, which were further speed vacuum-dried prior to LC–MS/MS analysis.

Samples (12 Fractions and the pooled sample) were resuspended in 5% formic acid in water and injected on an UltiMate 3000 RSLCnano System coupled to an Orbitrap Fusion Lumos Tribrid Mass Spectrometer (Thermo Scientific). Peptides were loaded on an Acclaim Pepmap trap column (Thermo Scientific #164750) prior analysis on a PepMap RSLC C18 analytical column (Thermo Scientific #ES903) and eluted on a 120 min linear gradient from 3 to 35% Buffer B (Buffer A: 0.1% formic acid in water, Buffer B: 0.08% formic acid in 80:20 acetonitrile: water (v:v)). Eluted peptides were then analyzed by the mass spectrometer operating in Synchronous Precursor Selection mode using a cycle time of 3s. MS1 were acquired at a resolution of 120000 with an AGC target of 100% and a maximum injection time of 50 ms. Peptides were then selected for MS2 fragmentation using CID with an isolation width of 0.7 Th, NCE of 35%, AGC of 100% and maximum injection time of 50 ms using the “rapid” scan rate. Up to 10 fragments were then selected for MS3 fragmentation using HCD with an isolation width of 3 Th, NCE of 65%, AGC of 200% and maximum injection time of 105 ms and spectra were acquired at a resolution of 50000. Dynamic exclusion was set to 60 s with a tolerance of +/- 10 ppm.

### Mass Spectrometry Data Analysis

Peptides were searched against Uniprot Swissprot Mouse containing isoforms (released on 2023/01/02) using MaxQuant (v2.2.0.0) ^64^. All parameters were left as default except for the addition of Deamidation (N, Q) and Phospho (S, T, Y) as variable modifications modification and the inclusion of MS3 TMT quantitation. Statistical analysis was carried out using Python (v3.9.0) and packages pandas (v1.3.3), numpy (v1.19.0), sklearn (v1.0), scipy (v1.7.1), rpy2 (v3.4.5), Plotnine (v0.7.1) and Plotly (v5.8.2) and R (v4.1.3) and the package *Limma* (3.50.1) ^65^. Sites identified as reverse, potential contaminant or with a localization confidence lower than 0.75 and sites quantified in less than 3 replicates in at least 1 condition were excluded. Missing values were then imputed using a gaussian distribution centered on the median with a downshift of 1.8 and width of 0.3 (relative to the standard deviation). Protein regulation was assessed using *limma* ^65^ and P-values were adjusted using Benjamini-Hochberg multiple hypothesis correction. Proteins were considered significantly regulated if their corrected P-value was smaller than 0.05 and their fold change was greater than 2 or smaller than 1/2. Data analysis was carried out using R (v4.2.3). Phospho-proteomic heatmaps were made using Morpheus (Broad Institute) https://software.broadinstitute.org/morpheus.

### Retroviral Transduction of T-IEL

Phoenix-Eco cells were maintained in DMEM (Gibco), 10% FBS, 1X Penicillin/Streptomycin, 1X pyruvate. 3x10^6^ cells were plated 24 hours prior to being transfected with 10 µg of either pMSCV-IRES-GFP or pMSCV-LAT-IRES-GFP combined with 10 µg of the packaging plasmid pCL-Eco using the calcium phosphate method. Briefly, the packaging plasmid was combined with the pCL-Eco plasmid, 50 µl CaCl2 and deionized water to a volume of 500 µl. 500 µl 2X HBS (140 mM NaCl, 1.5 mM Na2HPO4, 50 mM HEPES, pH 7.05) was added dropwise to the mixture while vortexing. The transfection mix was added dropwise to the Phoenix cells. Media was refreshed 12 – 16 hours after transfection. 48 hours after transfection virus was harvested and filtered through a 0.45 µm filter. Media was refreshed on the Phoenix cells and virus harvested at 72 hours post-transfection.

T-IEL were harvested and cultured in IL-15 following the previously described protocol. 48 hours after isolation, the cells were transduced with virus. Plates were coated with Retronectin (TakaraBio) and filtered supernatant of Phoenix cells producing virus loaded by two rounds of centrifugation. Cells were added to the plates containing the virus bound onto the Retronectin and spun briefly. Cells were incubated at 37°C overnight and the transduction process was repeated the following day. Transduced T-IEL were stimulated following the stimulation protocol described above.

### Single-cell RNA seq analyses

The generation of single-cell RNA sequencing (scRNA-seq) and TCR profiling data of murine and human IELs was previously described ^33^. Reanalysis of the data was performed using Seurat (v5.1.0) and in R (v4.4.2). Marker genes distinguishing natural αβ and γδ IELs from induced IELs subsets (i.e. CD8A, CD5, CD6, CD90, FCER1G) were used for cluster annotation. The scCustomize (v2.1.2) Clustered_DotPlot function was used to show the scaled expression level and percentage of cells expressing select genes for each T-IEL subset. ScRNA-seq data for porcine T-IEL was obtained from the USDA database (SCP1921)^43^, for mammary gland T-IEL from GSE279627^45^, for breast cancer and prostate cancer T cells from GSE195937^46^, and for human celiac disease IEL was obtained from GSE252762^6^. Bulk RNAseq data for mammary T-IEL were from GSE253989^45^, and for αβ innate like breast T cell thymic precursors and CD8SP from GSE195937^46^. Data processing and visualization were performed using Seurat (v4.0) in R (v4.4.2). For GSE252762, the dataset was loaded using readRDS(), and cells labeled as "Control" in the metadata were extracted via subset(). The SCP1921 dataset (Ileum_AllCells.h5seurat) was imported using the LoadH5Seurat() function from SeuratDisk. A curated list of marker genes (Figure 5D) was selected for visualization. Gene expression patterns across cell clusters were visualized using the custom_DotPlot() function from scCustomize, applying the "plasma" colormap from viridis to enhance contrast. Dot size represents the proportion of cells expressing a given gene, while color intensity reflects average expression levels. No additional filtering beyond subsetting for the control group and selected T cell subsets was applied. All dot plot visualizations were performed in R (v4.4.2) using scCustomize, ggplot2, viridis, and Seurat.

For Pseudotime ordering, scRNA-seq data (GSE122740) from IEL precursors ^34^ were analyzed using the RaceID pipeline. The RaceID object (IEL) was obtained from Dr. Patrice loaded into R, and a predefined pseudotemporal ordering of cells was obtained from the external dataset (pseudotime_IEL_cluster_5_3_1_4_2.csv). The pseudotemporal trajectory was extracted as a character vector, with cell types assigned after removing unique sample identifiers via regular expressions. A data frame was constructed to store pseudotime values and corresponding cell progression indices. Functionally relevant T cell signaling genes were selected for further analysis, and their expression levels were extracted from the normalized expression matrix (IEL@ndata). Expression trends were visualized using scatterplots with locally estimated scatterplot smoothing (LOESS) regression curves. All visualizations were performed in R (v4.4.2) using tidyverse, ggplot2, and RaceID.

### Proteomics of other T cell subsets

Regulatory T cells (nTregs) were FACS sorted from the spleens of Foxp3-YFP reporter mice based on YFP expression. Natural killer T (NKT) cells were isolated from the liver using mouse CD1d PBS-57 APC-labeled Tetramer (NIH Tetramer Facility) staining, followed by FACS sorting. Exhausted T cells were generated following the protocol described in ^47^. Natural T-IEL cells were isolated from the small intestinal epithelium and sorted into: Natural αβ T-IELs: TCRβ⁺CD103⁺CD8α⁺CD8β⁻CD4⁻ and Natural γδ T-IELs: TCRγδ⁺CD103⁺CD8α⁺CD8β⁻CD4⁻. Proteomic profiling was performed as previously established^4^, for quantification of proteins in each cell type.

## Data analysis

Flow cytometry data was analyzed using FlowJo Software (v10.9.0). Data analysis and statistical tests were carried out using GraphPad Prism (v10.0.0).

## Data availability

All data presented in this study are shown in the figures and extended data. The mass spectrometry proteomics data have been deposited to the ProteomeXchange Consortium via the PRIDE partner repository with the dataset identifier PXD063627.

