## Supplementary Figures 1-9 for "Conserved developmental rewiring of the TCR signalosome drives tolerance in innate-like lymphocytes"

### Developmental rewiring of the TCR signalosome in natural intestinal intraepithelial T lymphocytes

#### Supplemental data

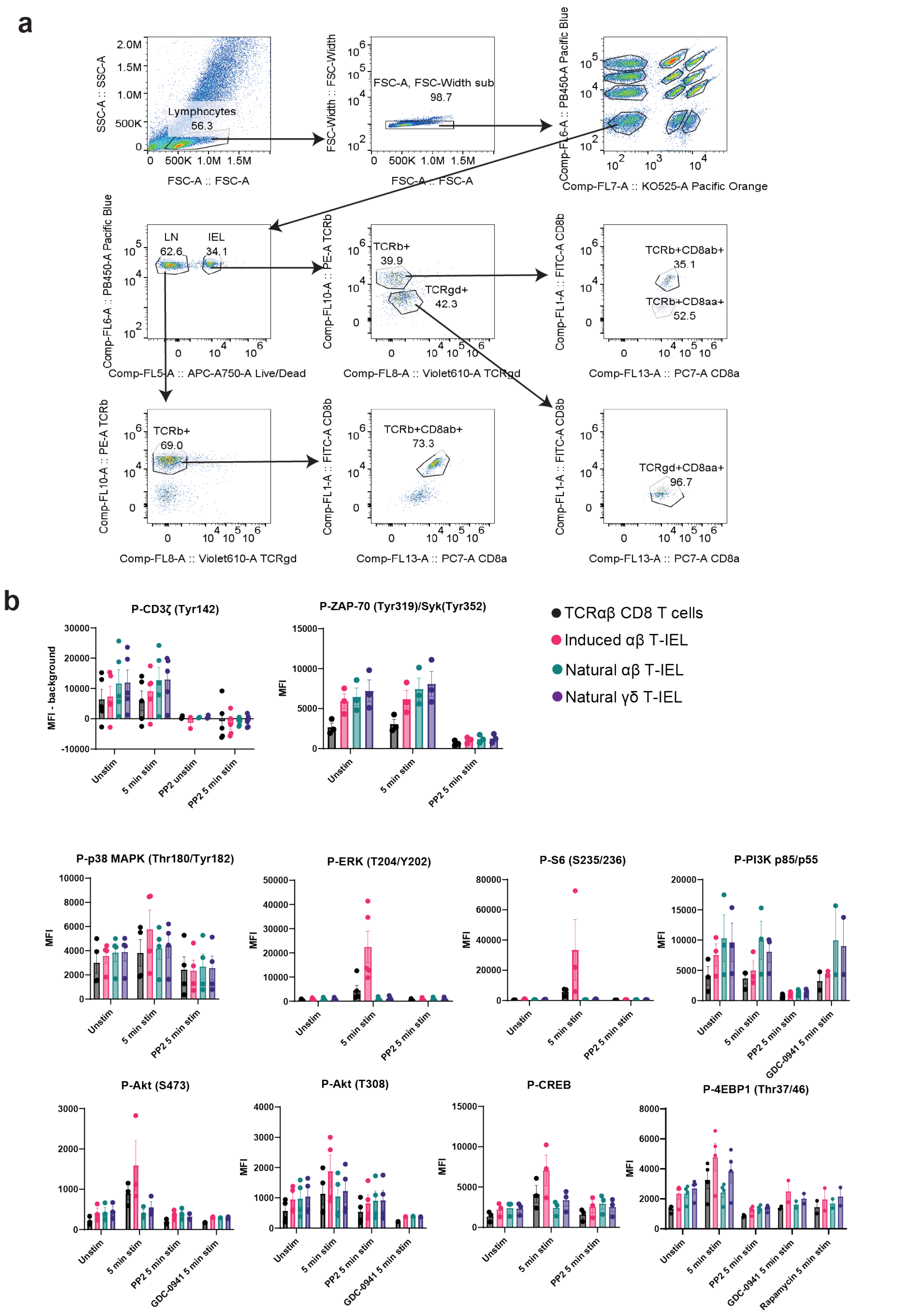


**Supplementary Figure 1: Gating Strategy for Phospho-Flow Barcoding and Inhibitor**

**Validation**. (a) Gating strategy for phospho-flow barcoding in TCR stimulation experiments. Lymphocytes were first gated by forward and side scatter (FSC/SSC) parameters. Single cells were identified using FSC-A vs. FSC-H, and individual stimulation conditions were distinguished based on distinct concentrations of Pacific Blue and Pacific Orange dyes. Conventional CD8α+ T cells were separated from T-IEL subsets using Near-IR amine-reactive dye staining. (b) Verification of antibody specificity and phosphorylation responses. Mean fluorescence intensity (MFI) of each phospho-protein in T-IEL subsets and conventional CD8α+ T cells after stimulation with anti-CD3 (30 μg/ml) crosslinked with anti-Hamster (5 μg/ml). Inhibitors were added 1 hour prior to stimulation, including PP2 (20 μM; Src family kinase inhibitor), Rapamycin (20 nM; mTOR inhibitor), and GDC-0941 (1 μM; PI3K inhibitor). Data represent mean ± SEM from three biological replicates.

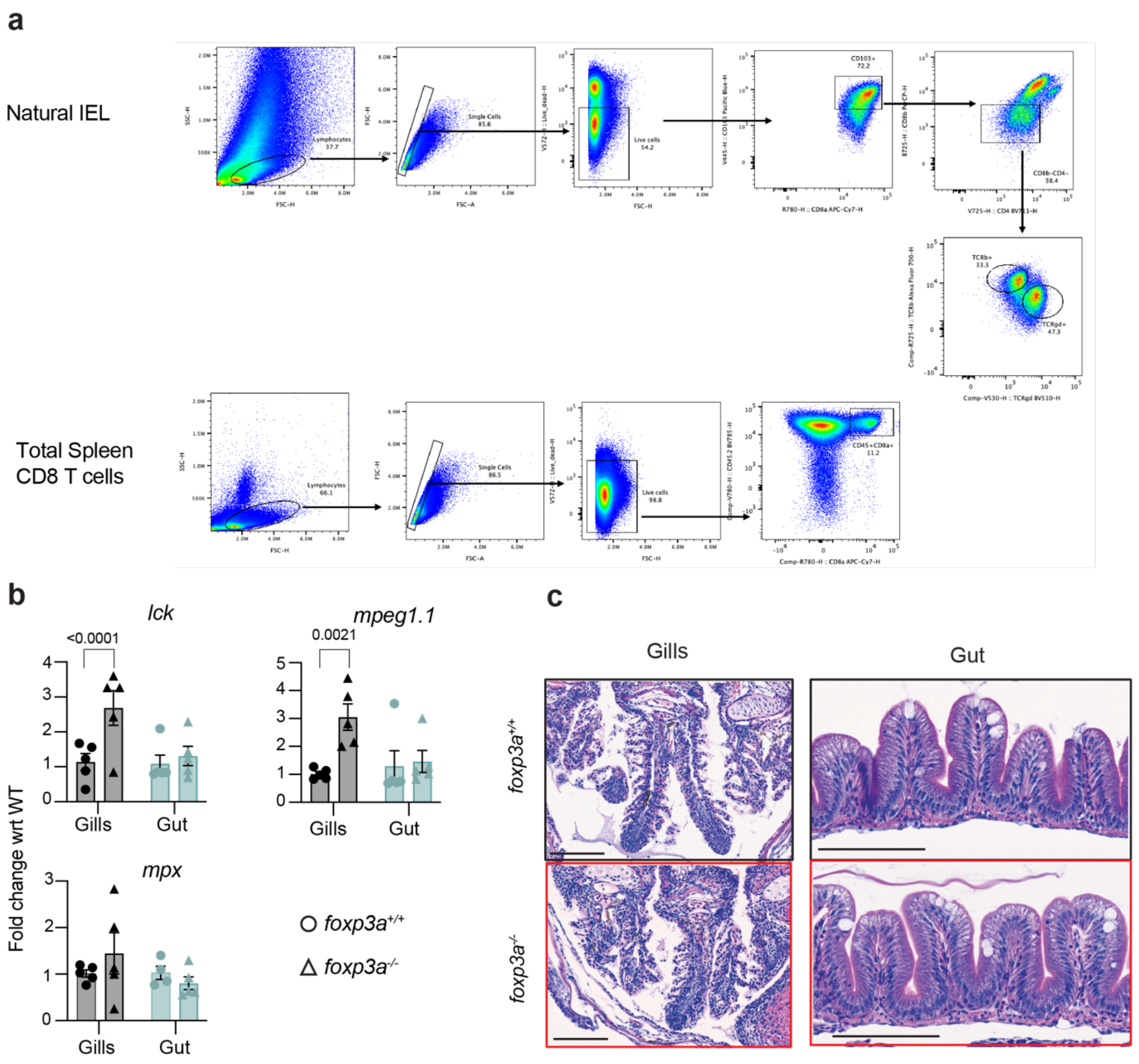


**Supplementary Figure 2: Gating strategy of conventional CD8α+ T cells and natural T-IEL subsets from FoxP3-DTR mice and analyses of *foxp3a$^{-/-}$* zebrafish. (a)** Flow cytometric gating strategy for isolating natural T-IEL subsets from the small intestine and conventional CD8+ T cells from the spleen. Lymphocytes were first gated based on forward and side scatter (FSC/SSC) parameters. Single cells were identified using FSC-A vs. FSC-H to exclude doublets. Live cells were gated based on viability dye exclusion, followed by CD8+ T cell selection in the spleen and natural T-IEL subset identification in the small intestine. **(b)** RT-qPCR analysis of immune cell marker gene expression in wild-type (WT) and *foxp3a$^{-/-}$* zebrafish (45 days post-fertilization, dpf), assessing immune activation in the absence of Foxp3. Data represents the mean ± SEM from five independent replicates. **(c)** Histological evaluation of inflammation in zebrafish gills and intestines. Representative H&E-stained

images of gills and gut from WT and foxp3a$^{-}$/$^{-}$ fish demonstrate tissue integrity and immune cell infiltration. Scale bar: 10 μm. Statistical significance for (a) and (c) was calculated using ANOVA; non-significant values are not shown.

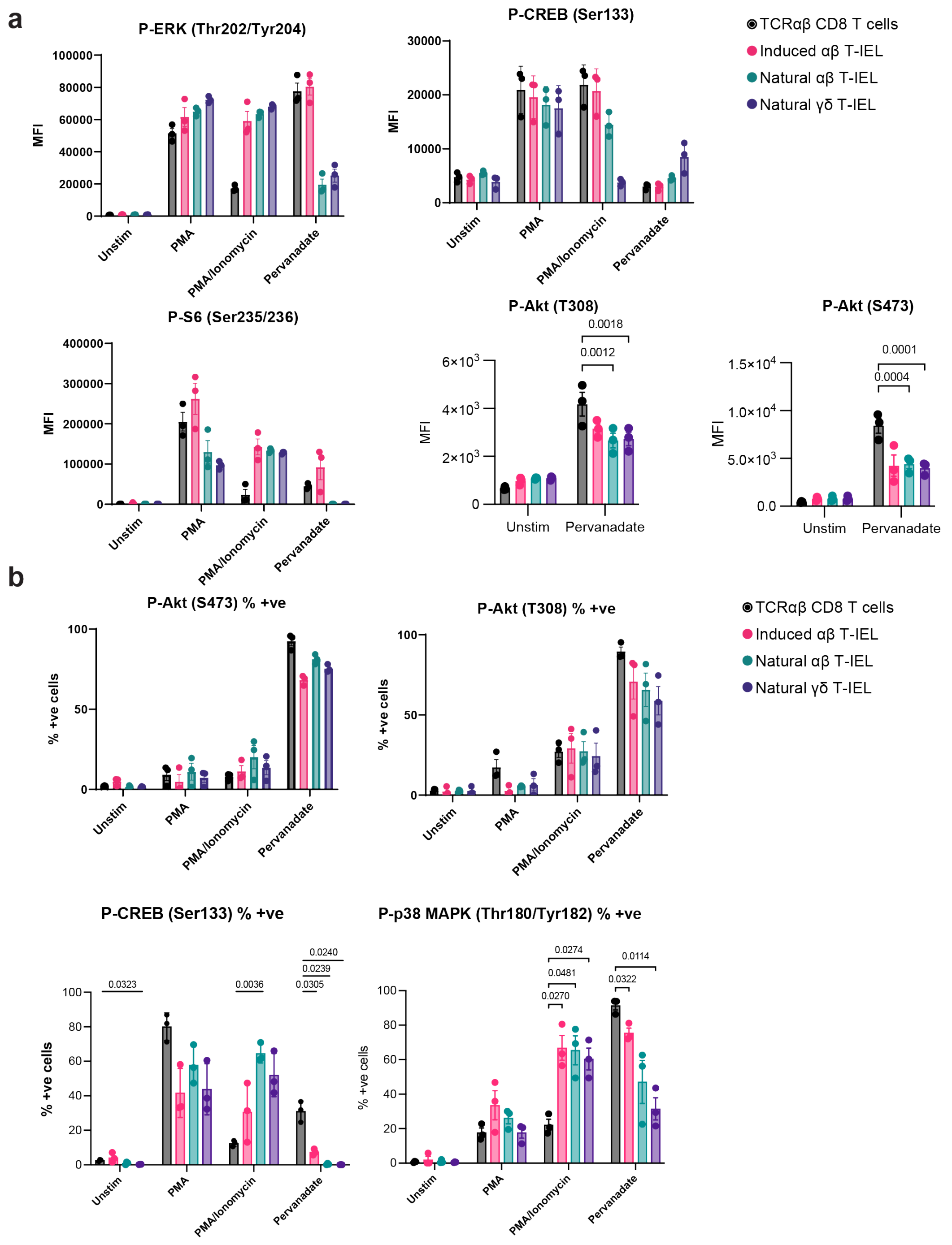


**Supplementary Figure 3: Bypassing proximal signaling permits functional TCR signaling responses in natural T-IELs**. Bar plots quantifying mean fluorescence intensity (MFI) of phospho-proteins and **(b)** bar plots depicting the percentage of phospho-protein-positive cells, upon stimulation with anti-CD3 (30 µg/ml) crosslinked with anti-Hamster (5 µg/ml), PMA (100 ng/ml), Ionomycin (1 µg/ml), or pervanadate (10 µM). Data represent mean ± SEM from three biological replicates.

Supp Figure 4

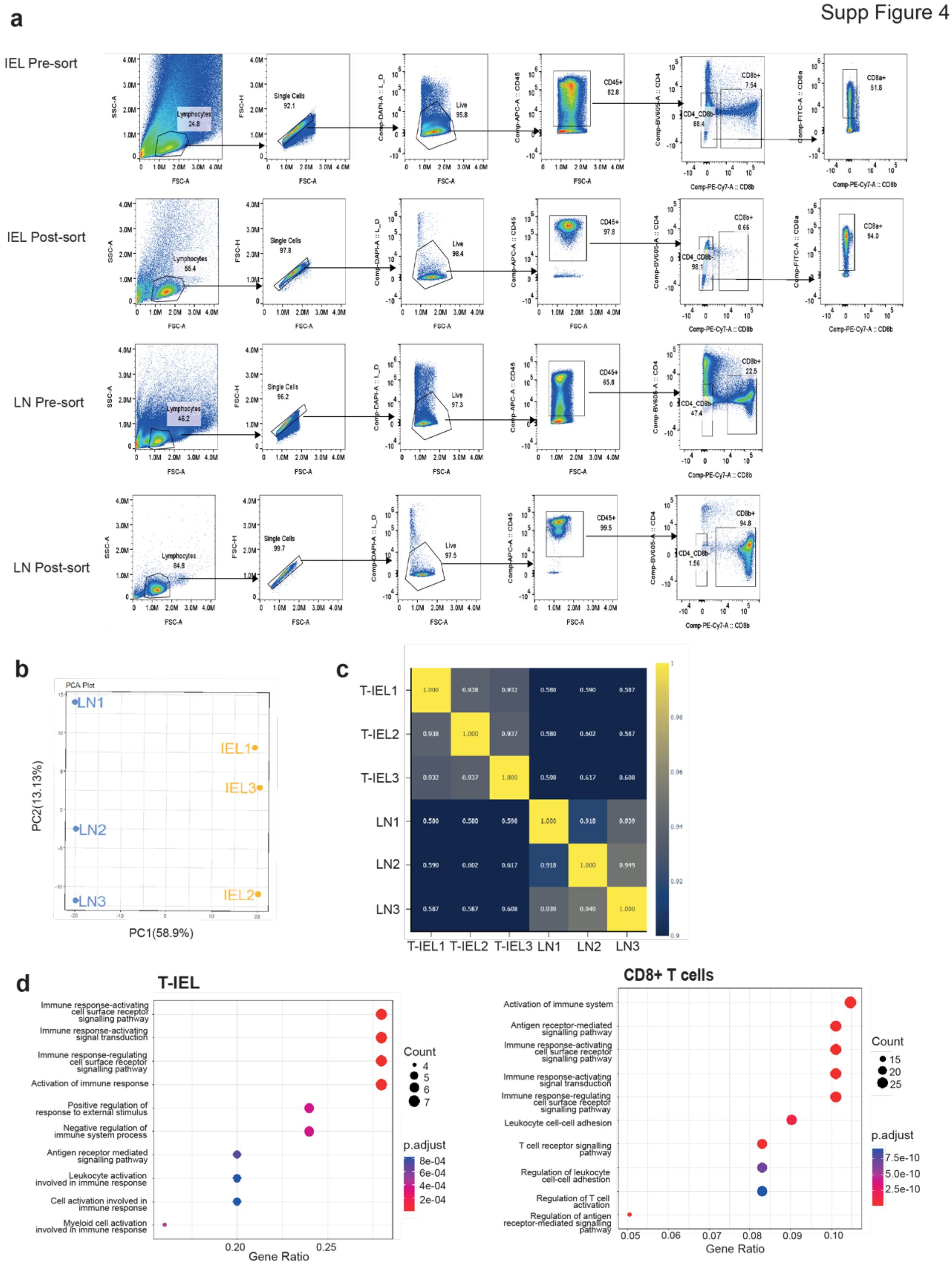


**Supplementary Figure 4: Distinct Phosphotyrosine signatures in natural T-IEL compared to conventional CD8+ T Cells**. **(a)** Representative flow cytometry profile showing the purity of conventional CD8 T cells sorted from lymph nodes (LN) and intestinal natural T-IEL populations following purification, prior to stimulation and phospho-proteomic analysis. **(b)** Principal component analysis (PCA) of natural T-IEL (IEL1-3) and conventional CD8 T cells

(LN1-3) phosphoproteomes, as shown in Figure 3f. **(c)** Correlation heatmap of T-IEL (IEL1-3) and conventional CD8+ T cell (LN1-3) phospho-proteomic samples. (d) Gene Ontology (GO) term enrichment analysis for proteins with significantly upregulated phospho-sites in natural T-IEL vs. conventional CD8+ T cells.

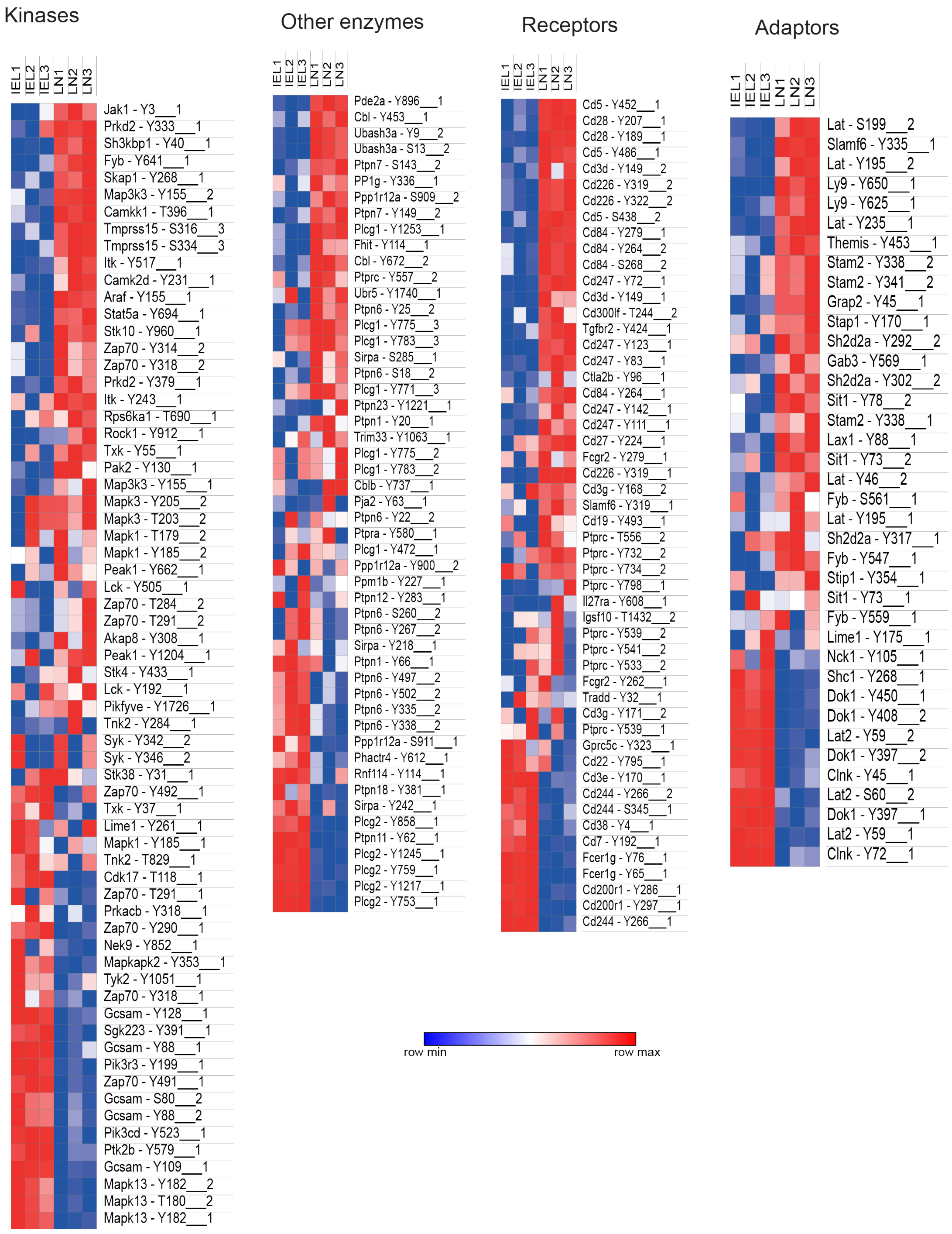


**Supplementary Figure 5: Phospho-proteomic profiling of natural T-IEL and conventional LN CD8 T cells**. Row-normalized heatmaps representing the median imputed phosphorylation intensities for the indicated phospho-sites in natural T-IEL (IEL1-3) and conventional CD8+ T cells (LN1-3) phosphoproteomes. Data were manually categorized into the main families of signaling proteins.

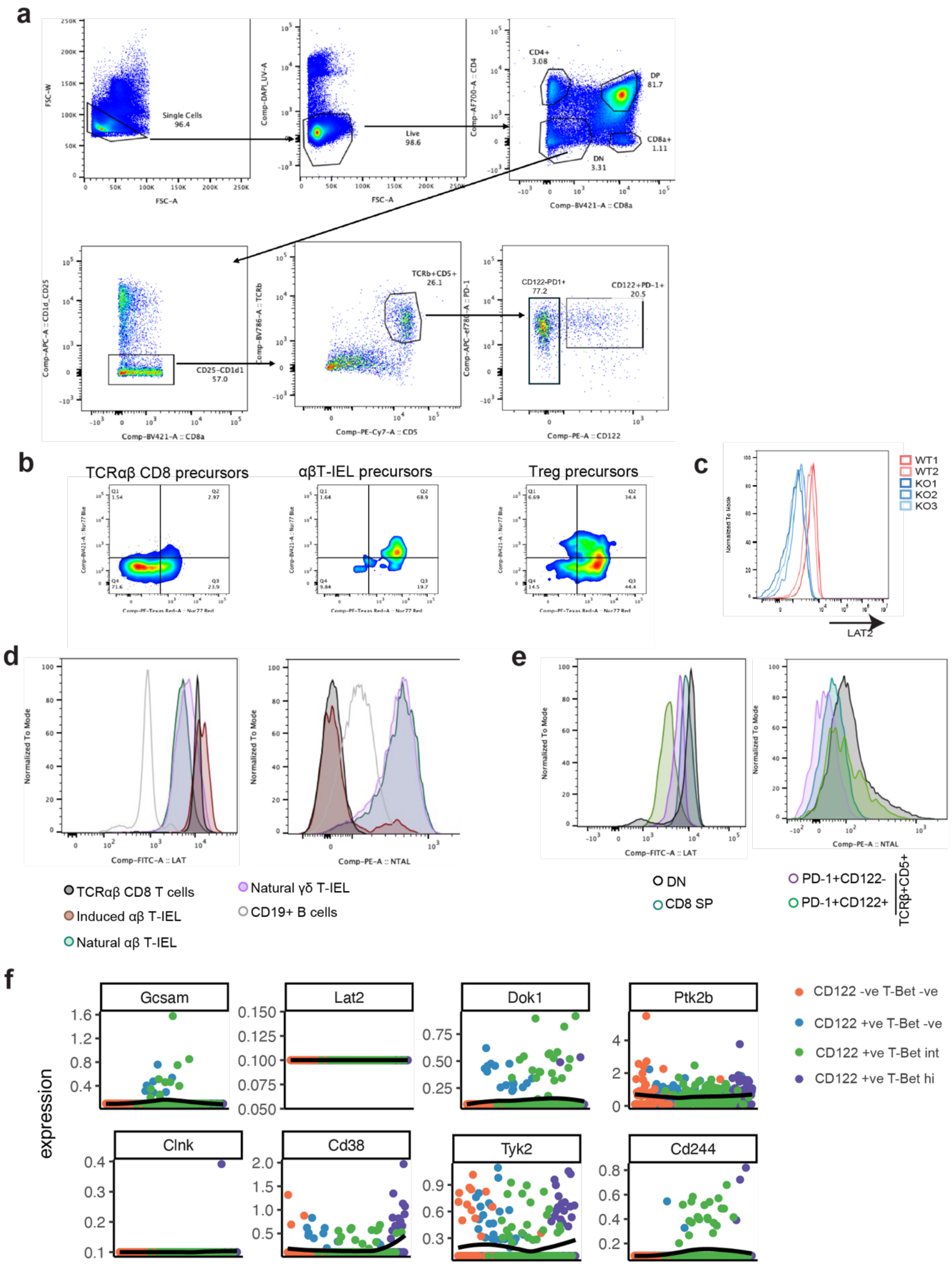


**Supplementary Figure 6: Analyses of thymic precursors of natural T-IEL**. **(a)** Gating strategy for identifying natural Treg precursors, natural αβ T-IEL precursors, and conventional CD8α+ T cell precursors in the thymus from Nur77-Tempo mice (for Fig. 4a). **(b)** Representative flow cytometry plots for Nur77-Blue and Nur77-Red fluorescence in thymic precursors of Tregs and natural αβ T-IEL and in CD8 single positive (SP) thymocytes. **(c)** Representative histogram validating specificity LAT2 antibody in wild-type T-IEL (red), with absence of LAT2 signal in LAT2 KO T-IEL (blue). **(d)** Representative histogram of LAT and LAT2 (NTAL) expression in mature intestinal T-IEL subsets compared to conventional CD8+ T cells. CD19+ B cells were used as negative and positive controls for LAT and LAT2 staining, respectively. **(e)** Representative histogram of LAT and LAT2 expression in thymocyte populations, including double-negative (DN) thymocytes ($CD4^{-}CD8^{-}$ $TCR\beta^{-}$), natural αβT-IEL precursors (gated as $CD4^{-}CD8^{-}TCR\beta^{+}CD5^{+}PD\text{-}1^{+}$ that are either $CD122^{-}$ or $CD122^{+}$), and conventional CD8+ T cell precursors (CD8 SP). **(e)** Expression trajectories of selected TCR signalosome genes along the predicted differentiation trajectory of natural αβ T-IEL precursors, analyzed from published single-cell RNA sequencing data[34].

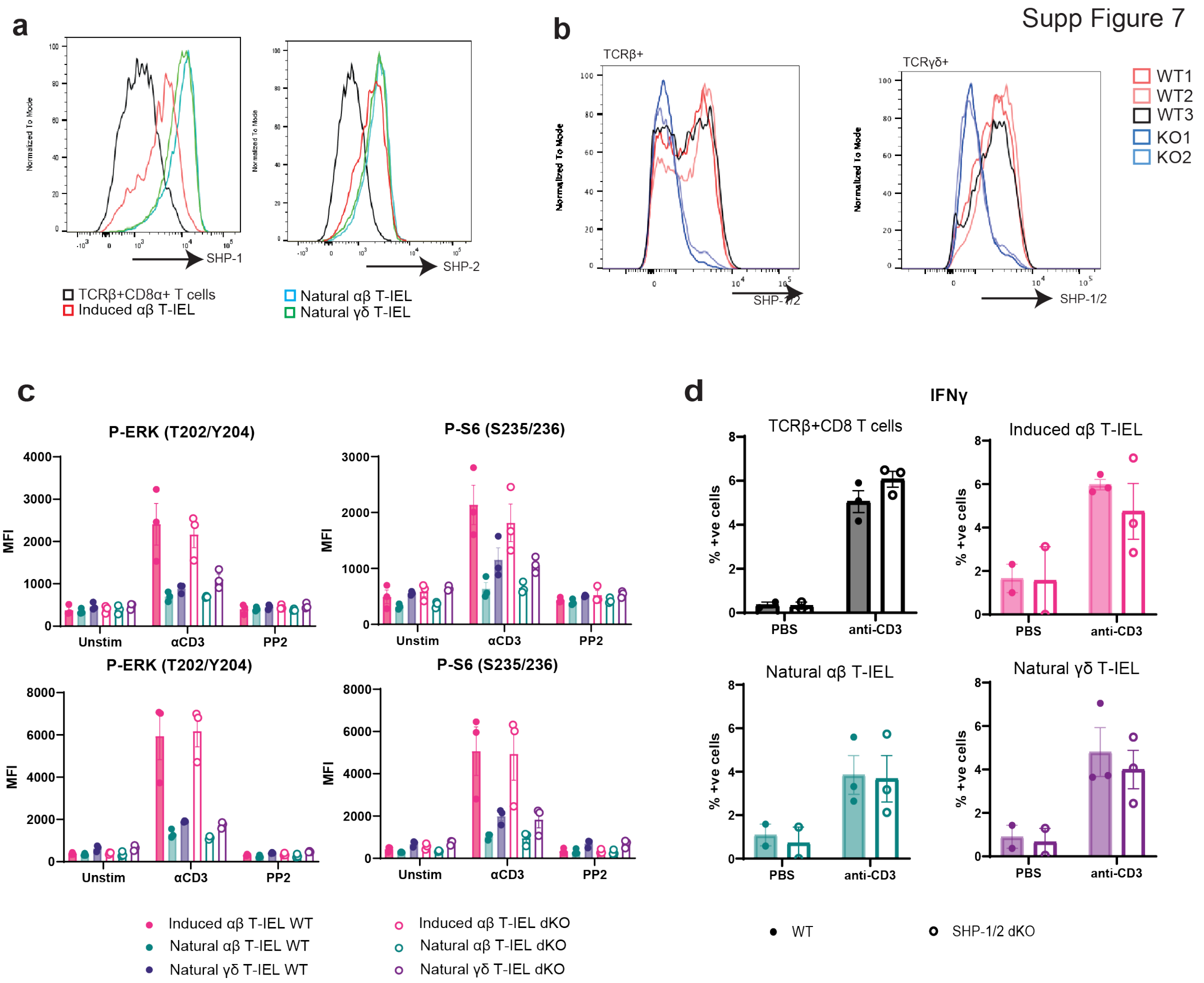


**Supplementary Figure 7: Functional Impact of SHP-1/2 Deficiency in Natural T-IEL**. **(a)** Histograms of SHP-1 and SHP-2 expression in T-IEL subsets and conventional CD8 T cells, confirming enrichment in natural T-IEL. **(b)** Histograms comparing SHP-1/2 expression in wild-type vs. IEL-specific SHP-1/2 knockout (GzmB-Cre x *Ptpn6*$^{fl/fl}$/*Ptpn11*$^{fl/fl}$ mice, SHP1/2 dKO) natural T-IEL subsets. **(c)** Bar graph of ERK and S6 phosphorylation MFI in wild-type vs. SHP-1/2 dKO T-IEL subsets upon in vitro anti-CD3 or pervanadate stimulation. Data represent mean ± SEM of three replicates. **(d)** Percentage of conventional CD8α+ T cells and T-IEL subsets positive for IFNγ production after i.p. injection of anti-CD3 (50 μg) or PBS vehicle control into WT and SHP-1/2 dKO mice.

Supp Figure 8

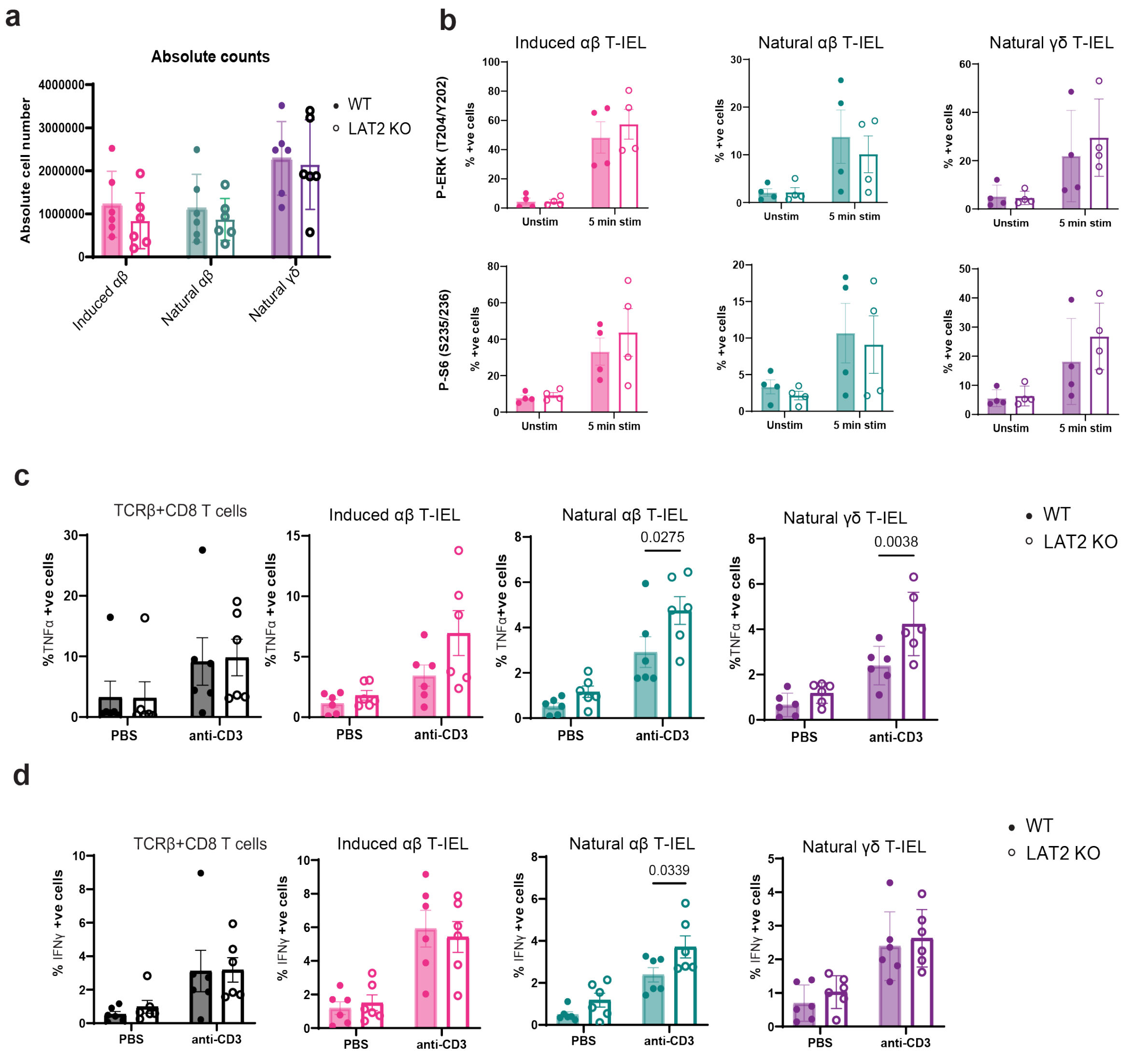


**Supplementary Figure 8: Impact of LAT2 Deficiency on Natural T-IEL signaling and Function**. **(a)** Absolute counts of natural T-IEL subsets in wild-type and LAT2 KO mice. (b) Percentage of T-IEL subsets positive for phosphorylation of ERK and S6 following *ex vivo* anti-CD3 stimulation, in wild-type (filled circles) and LAT2 KO (empty circles) T-IEL. Data represent mean ± SEM of four replicates. **(c-d)** Expression of TNF (c) and IFNγ (d) in conventional CD8 T cells from spleen and T-IEL subsets after i.p. injection of anti-CD3 (50 µg) or PBS vehicle control into LAT2 KO or WT littermate control mice. Data represent mean ± SEM of six biological replicates. Statistical significance for (b-d) was determined using ANOVA with Tukey's post hoc correction.

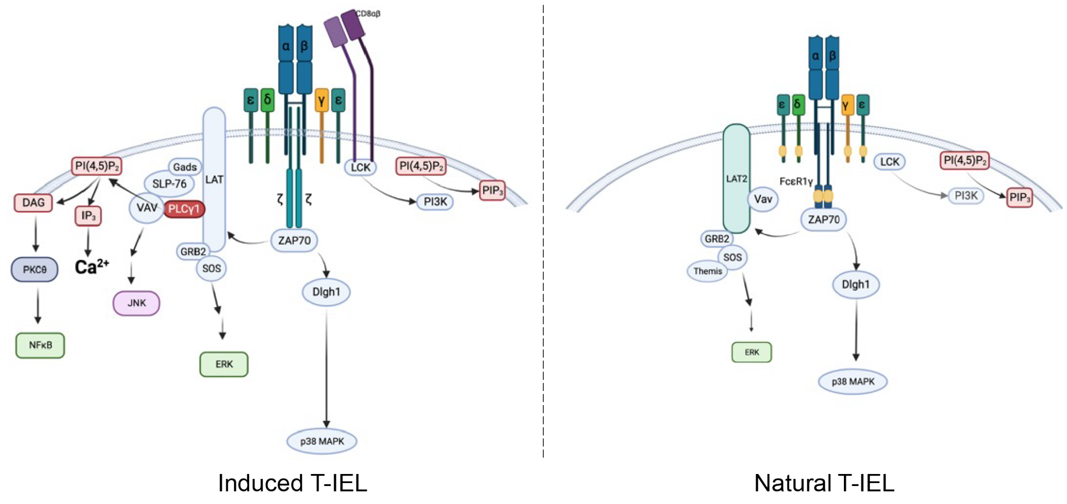


**Supplementary Figure 9.** Summary of the **RePrESS** (Rewiring of Proximal Elements of the TCR Signalosome for Suppression) model of suppression of TCR signaling in natural T-IEL, with proximal TCR signaling in conventional T cells shown as a comparison. Created in Biorender.com.
