## Supplemental Tables 1-3 for "Conserved developmental rewiring of the TCR signalosome drives tolerance in innate-like lymphocytes"

**Supplementary Table 1**

**Table 1:** List of genes included in each GO term category enriched in conventional CD8^+^ T cells.

| **GO Category** | **Genes included in the Category** |
| --- | --- |
| immune response-activating cell surface receptor signalling pathway | Cd28/Ubash3a/Cd226/Cd247/Skap1/Plcg1/Themis/Lax1/Prkd2/Ptpn6/Itk/Fyb/Zap70/Txk/Rftn1/Ptprc/Cblb/Cd19/Rab29/Usp9x/Syk/Mapk1/Lpxn/Cyld/Stap1/Ezr/Rbck1/Lck |
| immune response-activating signal transduction | Cd28/Ubash3a/Cd226/Cd247/Skap1/Plcg1/Themis/Lax1/Prkd2/Ptpn6/Itk/Fyb/Zap70/Txk/Rftn1/Ptprc/Cblb/Cd19/Rab29/Usp9x/Syk/Mapk1/Lpxn/Cyld/Stap1/Ezr/Rbck1/Lck |
| immune response-regulating cell surface receptor signalling pathway | Cd28/Ubash3a/Cd226/Cd247/Skap1/Plcg1/Themis/Lax1/Prkd2/Ptpn6/Itk/Fyb/Zap70/Txk/Rftn1/Ptprc/Cblb/Cd19/Rab29/Usp9x/Syk/Mapk1/Lpxn/Cyld/Stap1/Ezr/Rbck1/Lck |
| activation of immune response | Cd28/Ubash3a/Cd226/Cd247/Skap1/Plcg1/Themis/Lax1/Prkd2/Ptpn6/Itk/Fyb/Zap70/Txk/Rftn1/Ptprc/Cblb/Cd19/Rab29/Usp9x/Syk/Mapk1/Lpxn/Cyld/Stap1/Ezr/Rbck1/Lck/Hsp90aa1 |
| regulation of antigen receptor-mediated signalling pathway | Ubash3a/Cd226/Prkd2/Ptpn6/Ptprc/Cblb/Cd19/Rab29/Usp9x/Lpxn/Cyld/Stap1/Ezr/Lck |
| leukocyte cell-cell adhesion | Cd28/Cd5/Stat5a/Skap1/Lax1/Anxa1/Ptpn6/Pag1/Tgfbr2/Zap70/Stk10/Sirpa/Ptprc/Lgals1/Cblb/Rock1/Cd27/Syk/Il27ra/Cyld/Coro1a/Dock8/Lck/Hsp90aa1/Capn1 |
| regulation of leukocyte cell-cell adhesion | Cd28/Cd5/Stat5a/Skap1/Lax1/Anxa1/Ptpn6/Pag1/Tgfbr2/Zap70/Sirpa/Ptprc/Lgals1/Cblb/Cd27/Syk/Il27ra/Cyld/Coro1a/Dock8/Lck/Hsp90aa1/Capn1 |
| regulation of T cell activation | Cd28/Cd5/Lat/Stat5a/Lax1/Anxa1/Sit1/Ptpn6/Pag1/Tgfbr2/Zap70/Sirpa/Ptprc/Lgals1/Cblb/Cd27/Syk/Il27ra/Cyld/Coro1a/Dock8/Lck/Hsp90aa1 |
| lymphocyte differentiation | Cd28/Ly9/Slamf6/Stat5a/Dock2/Themis/Cd3d/Anxa1/Ptpn6/Itk/Dock11/Tgfbr2/Zap70/Txk/Nhej1/Ptprc/Lgals1/Cd19/Cd27/Syk/Cyld/Lck/Cd3g/Ikzf3/Hsp90aa1 |
| positive regulation of cell-cell adhesion | Cd28/Cd5/Stat5a/Skap1/Anxa1/Tgfbr2/Zap70/Sirpa/Ptprc/Jak1/Lgals1/Cd27/Syk/Il27ra/Cyld/Coro1a/Dock8/Ptpn23/Lck/Hsp90aa1/Capn1 |
| T cell selection | Cd28/Ly9/Slamf6/Dock2/Themis/Cd3d/Zap70/Ptprc/Syk/Cyld/Cd3g |
| regulation of cell-cell adhesion | Cd28/Cd5/Stat5a/Skap1/Lax1/Anxa1/Ptpn6/Pag1/Tgfbr2/Zap70/Sirpa/Ptprc/Jak1/Lgals1/Cblb/Cd27/Syk/Il27ra/Cyld/Coro1a/Dock8/Ptpn23/Lck/Hsp90aa1/Capn1 |
| positive regulation of leukocyte cell-cell adhesion | Cd28/Cd5/Stat5a/Skap1/Anxa1/Tgfbr2/Zap70/Sirpa/Ptprc/Lgals1/Cd27/Syk/Il27ra/Cyld/Coro1a/Dock8/Lck/Hsp90aa1/Capn1 |
| leukocyte proliferation | Cd28/Slamf6/Stat5a/Dock2/Anxa1/Ptpn6/Sh2d2a/Tgfbr2/Zap70/Ptprc/Cnn2/Cblb/Cd19/Cd27/Syk/Il27ra/Mapk1/Coro1a/Rassf5/Dock8/Mapk3/Ikzf3 |
| regulation of T cell receptor signalling pathway | Ubash3a/Cd226/Prkd2/Ptpn6/Cblb/Rab29/Usp9x/Cyld/Ezr/Lck |
| T cell differentiation | Cd28/Ly9/Slamf6/Stat5a/Dock2/Themis/Cd3d/Anxa1/Itk/Tgfbr2/Zap70/Txk/Nhej1/Ptprc/Cd27/Syk/Cyld/Lck/Cd3g/Hsp90aa1 |
| positive T cell selection | Ly9/Slamf6/Dock2/Themis/Cd3d/Zap70/Ptprc/Cyld/Cd3g |
| positive regulation of T cell activation | Cd28/Cd5/Stat5a/Anxa1/Tgfbr2/Zap70/Sirpa/Ptprc/Lgals1/Cd27/Syk/Il27ra/Cyld/Coro1a/Dock8/Lck/Hsp90aa1 |
| lymphocyte proliferation | Cd28/Slamf6/Stat5a/Dock2/Anxa1/Ptpn6/Sh2d2a/Tgfbr2/Zap70/Ptprc/Cblb/Cd19/Cd27/Syk/Il27ra/Coro1a/Rassf5/Dock8/Ikzf3 |
| mononuclear cell proliferation | Cd28/Slamf6/Stat5a/Dock2/Anxa1/Ptpn6/Sh2d2a/Tgfbr2/Zap70/Ptprc/Cblb/Cd19/Cd27/Syk/Il27ra/Coro1a/Rassf5/Dock8/Ikzf3 |

**Supplementary Table 2**

**Table 2**: List of genes included in each GO term category enriched in CD8a^+^ IEL.

| **GO Category** | **Genes included in the Category** |
| --- | --- |
| immune response-activating cell surface receptor signalling pathway | Fcer1g/Lat2/Plcg2/Gcsam/Masp1/Zap70/Cd38 |
| immune response-activating signal transduction | Fcer1g/Lat2/Plcg2/Gcsam/Masp1/Zap70/Cd38 |
| immune response-regulating cell surface receptor signalling pathway | Fcer1g/Lat2/Plcg2/Gcsam/Masp1/Zap70/Cd38 |
| activation of immune response | Fcer1g/Lat2/Plcg2/Gcsam/Masp1/Zap70/Cd38 |
| positive regulation of response to external stimulus | Fcer1g/Clnk/Plcg2/Masp1/Ptk2b/Fbn1 |
| negative regulation of immune system process | Fcer1g/Cd200r1/Clnk/Gcsam/Masp1/Fbn1 |
| myeloid cell activation involved in immune response | Fcer1g/Lat2/Clnk/Plcg2 |
| antigen receptor-mediated signalling pathway | Lat2/Plcg2/Gcsam/Zap70/Cd38 |
| leukocyte activation involved in immune response | Fcer1g/Lat2/Clnk/Plcg2/Ptk2b |
| cell activation involved in immune response | Fcer1g/Lat2/Clnk/Plcg2/Ptk2b |
| regulation of interleukin-6 production | Fcer1g/Cd200r1/Mapk13/Plcg2 |
| interleukin-6 production | Fcer1g/Cd200r1/Mapk13/Plcg2 |
| B cell receptor signalling pathway | Lat2/Plcg2/Gcsam/Cd38 |
| calcium-mediated signalling | Lat2/Plcg2/Zap70/Ptk2b |
| regulation of lymphocyte migration | Cd200r1/Gcsam/Ptk2b |
| mast cell degranulation | Fcer1g/Lat2/Clnk |
| B cell activation | Lat2/Plcg2/Pik3cd/Ptk2b/Cd38 |
| mast cell mediated immunity | Fcer1g/Lat2/Clnk |

**Supplementary Table 3**

**Table X**: List of primers used in zebrafish study

| **Oligo ID** | **Sequence** |
| --- | --- |
| actb2-F | GCCTGACGGACAGGTCAT |
| actb2-R | ACCGCAAGATTCCATACCC |
| lck-F | GTTATGGGACATGGCAACG |
| lck-R | CAAAAGAGCCGCAGTTCC |
| mpeg1.1-F | TGTTACAGCACGGGTTCAAG |
| mpeg1.1-R | AATGGCGTCAGCGATTTC |
| mpx-F | TTCCAGAAAATCCGAGATGG |
| mpx-R | ACAGAGGCCAGAGCTGTTTT |
